# pamiR: INVESTIGATING PLANT CELLS ONE ORGANELLE AT A TIME

**DOI:** 10.64898/2026.03.12.711057

**Authors:** Benjamin Brandt, Anna I Pratt, Carina Engstler, Deborah Schwarz, Dominik Schneider, Felix Hauser, Chase L Lewis, Chance M Lewis, Serena Schwenkert, Rainer Schwacke, Hans-Henning Kunz

**Affiliations:** Plant Biochemistry and Physiology, LMU München, Großhaderner Straße 2-4, 82152 Martinsried-Planegg, Germany; Plant Physiology, School of Biological Sciences, Washington State University, PO Box 644236, Pullman, WA 99164-4236, USA; Compact Plants Phenomics Center, Washington State University, PO Box 646340, Pullman, WA 99164-6340, USA; Division of Biological Sciences, Cell and Developmental Biology Section, University of California San Diego, La Jolla, California 92093-0116; Plant Molecular Biology, LMU München, Großhaderner Straße 2-4, 82152 Martinsried-Planegg, Germany; Mass Spectrometry of Biomolecules (MSBioLMU), LMU München, Großhaderner Straße 2-4, 82152 Martinsried-Planegg, Germany; IBG-4 Bioinformatics, Forschungszentrum Jülich, Wilhelm-Johnen-Straße, 52428 Jülich, Germany

## Abstract

Plant cell biochemistry is highly compartmentalized. Hence, forward genetic screening in an organelle-specific manner holds great potential to reveal novel loci controlling cell physiology. However, conventional forward genetics is a) limited by functional redundancy that can mask phenotypes in plant gene families and b) lacks organelle specificity. Here, we present the plastid-focused artificial microRNA (*pamiR*) resource designed to silence individual nuclear genes or co-targetable gene-family subsets encoding the organelle’s proteome. The library contains 2,285 artificial microRNAs directed against 2,202 genes selected from a plastid-enriched, proteomics-supported target set. The constructs are expressed by a constitutive UBQ10 promoter in a binary vector carrying the FAST seed-fluorescence marker, allowing transformant selection before germination. Two proof-of-concept screens, one using photosynthetic phenotyping and one using paclobutrazol, a gibberellin biosynthesis inhibitor, recovered lines with phenotypes previously associated with simultaneous perturbation of multiple members of redundant plastid gene families such as light-harvesting complex b1 (LHCB1), K^+^ efflux antiporter (KEA), and 9-cis-epoxycarotenoid dioxygenase (NCED). We experimentally confirmed specific target depletion for selected candidates, including paralogue-resolved LHCB1 depletion by mass spectrometry, and provide predicted off-target annotations for all 2,285 guides. Untargeted LHCB family members remained unchanged, and across the proteins included in the proteome-wide differential-abundance analysis, no additional significantly decreased protein was detected. KEA1 immunoreactive signal also fell below the immunoblot detection limit in *pamiRkea1/2* and an independent *KEA1/KEA2* amiRNA line. Together, these data establish *pamiR* as a plastid-specific screening resource and provide a framework for prioritizing and independently validating candidate phenotypes.

**One-sentence summary:** The *pamiR* library enables family-level silencing of nuclear genes encoding plastid proteins and provides per-construct target and predicted off-target annotations for organelle-focused forward genetic screens.

## Introduction

In 2000, *Arabidopsis thaliana*, the most studied plant in the world, was among the first eukaryotes with a sequenced genome (The Arabidopsis Genome, 2000). The advent of publicly available loss-of-function mutants, primarily using the Transfer-DNA (T-DNA) genome insertion approach (McKinney et al., 1995), made gene discovery a real success story with tremendous functional insights not limited to plant science (Provart et al., 2016). In addition, translational research has improved many crop plant species and spurred a slew of biotechnological innovations (Yaschenko et al., 2024). However, a little over 25 years after the release of the genome sequence, the proportion of experimentally characterized genes in *A. thaliana* remains below 33% (Hansen et al., 2018; Depuydt and Vandepoele, 2021). Moreover, ongoing discoveries of novel or additional functions for already “characterized” genes draw a more complex picture and emphasize the need for innovative tools to complete our functional understanding of the plant genome, which is the basis for applied plant biotechnology.

Forward genetics is a particularly suitable strategy to unveil gene functions, since the alteration of a known and valuable genetic trait is the starting point of such investigations (Page and Grossniklaus, 2002). Several molecular means enable forward genetic screens e.g., Ethyl Methanesulphonate (EMS), fast neutron, and transposon mutagenesis. However, since simultaneous loss-of-function mutations within two or more members of a gene families are statistically highly unlikely, these approaches lack the power to overcome functional genetic redundancy (FGR). FGR is prevalent within gene families frequently found in plant genomes. Here, a gene loss is compensated by another family member so that informative phenotypes only emerge in higher-order mutants (Iohannes and Jackson, 2023).

Two orthogonal strategies can overcome FGR: multi-target genome-scale CRISPR libraries (Hu et al., 2023) and genome-wide family-specific artificial microRNAs (amiRNAs (Schwab et al., 2006; Hauser et al., 2013)). While CRISPR/Cas-mediated editing affects gene function by modifying the genome sequence, amiRNAs silence genes post transcriptionally. Both approaches have their pros and cons but were successfully employed to unveil new gene functions (Xie et al., 2021; Hu et al., 2023). Recently, the Multi Targeted AmiRNA Cell type–specific Transportome-scale (mTACT) library was introduced, which enables cell type-specific silencing of transporter genes (Anfang et al., 2025). However, none of these tools consider distinct subcellular localizations of gene family products. Thanks to genome duplications and subsequent sequence adjustments this phenomenon has profoundly shaped plant cell biochemistry and physiology. A premier example is the K^+^ efflux antiporter (KEA) family with its six members that reside in two different plastid membranes, the Golgi, the trans-Golgi network, and the prevacuolar compartment/multivesicular bodies (Höhner et al., 2016; Zhu et al., 2018). Since the current genomic resources do not account for this inherent feature, often pleiotropic hard to interpret mutant phenotypes emerge from the simultaneous suppression of gene functions within distinct cellular compartments.

To resolve the issue, we built the first organelle-specific, plastid-focused amiRNA (*pamiR*) tool that bypasses FGR. The distribution of proteins across the organelles of *A. thaliana* has been reasonably well established (Fig. 1A; (Hooper et al., 2017)). We focused on the plastid, the central hub for plant productivity and stress tolerance, that harbors photosynthesis and highly specialized pathways e.g., to produce phytohormones or their precursors (Kunz et al., 2024).

**Figure 1.**
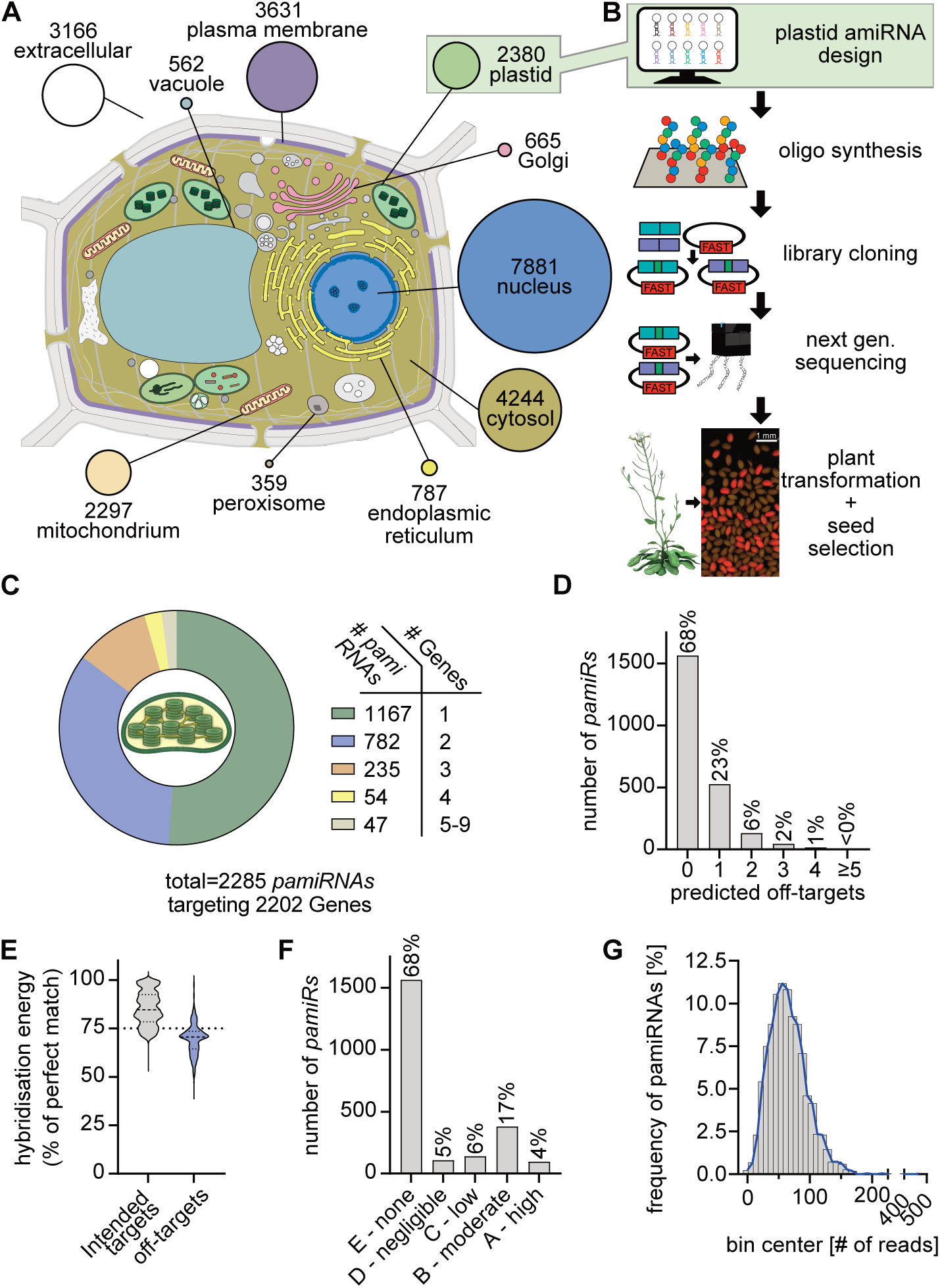
Design, construction, quality control, and predicted off-target landscape of the plastid artificial microRNA (*pamiR*) library. A) Contextual subcellular-proteome sizes based on SUBA/SUBAcon consensus assignments. B) Workflow from plastid-target selection and *pamiR* design through oligonucleotide synthesis, pooled cloning, next-generation sequencing (NGS), plant transformation, and FAST-based fluorescent seed selection. The seed-fluorescence image is illustrative and is not intended to represent T_1_ transformation frequency or Mendelian segregation. C) Number of intended target genes per *pamiR* D) Number of additional WMD3-predicted transcript matches per guide. E) Relative hybridization energy of recovered intended-target and additional predicted alignments. F) Per-guide A-E prioritization class based on the highest-impact additional predicted alignment. Class E denotes no additional transcript returned under the stated WMD3 settings; classes A-D prioritize returned alignments using mismatch position and relative hybridization energy as defined in Materials and Methods. These classes are prediction/prioritization categories and are not experimentally calibrated probabilities of off-target silencing. G) NGS-based representation of the final plasmid pool.

Three questions structure this study: 1) How was the plastid target set assembled, and how strong is the localization evidence underlying it? 2) How specifically do the designed guides act on their intended targets, and how is the residual predicted off-target risk reported? 3) Can the *pamiR* library recover interpretable phenotypes from single transformation events in redundant gene families? Besides other experiments, we address these questions through a proteomics-supported target-space curation, a library-wide database query with per-construct predicted off-target annotation, protein-level validation in two unrelated gene families, and two proof-of-concept screens. Because forward genetics screens initially produce candidates rather than definitive causal relationships, we also define the independent validation steps required before a *pamiR* phenotype can be interpreted mechanistically.

## Results and Discussion

### A proteomics-supported, plastid-enriched target set of nuclear encoded proteins

For an initial compartment proteome overview, 26,882 Araport11 entries were queried in the SUBcellular location database for Arabidopsis proteins (SUBA) and filtered to retain proteins with a single SUBA consensus assignment, leaving 25,972 entries. The compartment protein numbers (Fig. 1A) are therefore mutually exclusive consensus assignments that sum to 25,972. SUBA integrates experimental localization evidence with computational prediction, so the evidence density is not uniform across proteins or compartments (Hooper et al., 2014; Hooper et al., 2017). Therefore, these values primarily illustrate the scale of the major subcellular proteomes.

The target space for nuclear genes encoding plastid proteins was assembled by surveying independent published chloroplast and plastid-membrane proteomic datasets (Sun et al., 2008; Ferro et al., 2010; Simm et al., 2013; Tomizioli et al., 2014; Hooper et al., 2017; Lundquist et al., 2017; Trentmann et al., 2020) and was subsequently refined using SUBA/MMAP localization information and additional literature. Because the initial candidate set was derived largely from proteomic studies, and because SUBA/SUBAcon integrates experimental localization evidence with computational predictions, this strategy strongly enriched the library for proteins with direct experimental support for plastid localization. We therefore did not perform an additional orthogonal localization experiment for every individual candidate. Following this curation and subsequent *pamiR* design, the final library comprised 2,285 amiRNAs targeting 2,202 unique genes (Figs. 1A-C and S1A; Supplementary Dataset S1).

Proteins with dual localization were retained, including proteins targeted to both plastids and mitochondria, as frequently observed for factors involved in tRNA aminoacylation and organellar gene expression (Brandt et al., 2026). In addition, localization may be context-or tissue-dependent, and assignment of a protein to the plastid does not exclude additional localization under other conditions.

### Family-targeting amiRNA design and library coverage

To prepare for our intended plastid specific forward genetic approach (Fig. 1B), the 21-nt amiRNA guides were designed using the well-established Web MicroRNA Designer (WMD) strategy previously employed in the construction of the PHANTOM collection (Schwab et al., 2006; Ossowski et al., 2008; Hauser et al., 2013). The aim was not to define gene families by a fixed global sequence-identity threshold. Instead, family members were grouped operationally when one guide could satisfy the WMD target-recognition criteria for all intended transcripts with the same organellar destination; families were split into smaller co-targetable subsets when no single guide covered the full group. These guide–gene relationships are visualized in Supplementary Fig. S1B, which illustrates both whole-family targeting and targeting of co-targetable subsets.

For intended targets, the design rules constrained complementarity in the 5-prime/central region of the guide, including the cleavage-centered positions 10/11, and required sufficient predicted duplex stability (Schwab et al., 2006; Ossowski et al., 2008). The full parameter set is given in the Materials and Methods section.

The final collection contains 2,285 amiRNAs directed against 2,202 unique intended genes. Of the guides, 1,167 (51.1%) have one intended target, 782 (34.2%) have two, 235 (10.3%) have three, 54 (2.4%) have four, and 47 (2.1%) have five to nine, giving a mean of 1.72 intended genes per amiRNA (median 1; range 1-9; Fig. 1C). Viewed from the gene perspective, 1,764 genes are represented by one construct and 438 by two or more constructs. These additional constructs can provide useful independent reagents, but they are not automatically equivalent because guide sequence, target site, and insertion event differ.

### A library-wide audit defines the predicted off-target space

Sequence and duplex features used by WMD are experimentally supported determinants of plant miRNA/amiRNA targeting. Direct amiRNA targets can be predicted with high specificity from established plant miRNA pairing rules (Schwab et al., 2006; Ossowski et al., 2008). Protein-based testing further identified target pairing and relative hybridization energy as important determinants of silencing efficacy, with optimal amiRNAs generally exceeding 80% of the perfect-match hybridization energy and often causing strong protein depletion (Li et al., 2013). These findings provide the empirical basis for prioritizing predicted off-targets according to mismatch position and duplex energy.

We therefore queried all 2,285 guides with WMD3 against Araport11, the most recent Arabidopsis cDNA reference available in WMD3 (release 201606), using the search settings specified in Materials and Methods. The resulting alignments represent predicted transcript interactions under these rules and complement, rather than replace, experimental validation (Fig. 1D-F; Supplementary Datasets S1 and S2). Across the complete query, 3,834 intended-target alignments and 1,028 additional predicted alignments were recovered (Supplementary Dataset S2). Most guides showed only a limited number of additional predicted interactions. Of the 2,285 *pamiRs*, 68% returned no additional transcripts and 23% returned one, whereas only approximately 9% returned more than one predicted off-target (Fig. 1D). Intended-target duplexes showed a substantially higher median relative hybridization energy than additional predicted alignments (84.7% versus 70.5%; Fig. 1E; Supplementary Dataset S2). In addition, only 2 of 3,834 intended alignments (0.05%) contained a mismatch at guide positions 10 or 11, compared with 210 of 1,028 additional predicted alignments (20.4%). Similarly, only 1 intended alignment (0.03%) contained at least two mismatches within positions 2–12, compared with 350 additional predicted alignments (34.0%; Supplementary Dataset S2).

To make the predicted off-target information directly useful and easily accessible for candidate evaluation, we classified each additional alignment into an ordered A–D risk category based on WMD sequence/duplex criteria and experimentally supported determinants of plant amiRNA silencing efficiency (Schwab et al., 2006; Li et al., 2013). The classes rank predicted amiRNA-mediated silencing potential from A (high) > B (moderate) > C (low) > D (negligible), whereas class E denotes *pamiRs* for which no off-target was returned under the stated WMD3 search settings. Classification is hierarchical, with cleavage-centered and 5′/central pairing constraints considered before overall duplex stability, reflecting their importance in the WMD target-recognition rules and experimental studies of plant miRNA targeting. Each *pamiR* is assigned the class of its highest impact predicted off-target. The underlying criteria, mechanistic rationale, and distribution of alignments and *pamiRs* across the classes are summarized in Supplementary Table S4, while *pamiR*-specific assignments are provided in Supplementary Dataset S1. Under this classification, 1,564 guides (68.4%) were class E, 246 (10.8%) had only class C or D interactions, 380 (16.6%) were assigned to class B, and only 95 guides (4.2%; 98 alignments) contained at least one class-A predicted off-target (Fig. 1F). Class A requires no mismatch at positions 10/11, at most one mismatch within positions 2–12, and ≥75% relative hybridization energy. This threshold was deliberately set below the >80% relative energy associated with optimal amiRNA activity in protein-based experiments (Li et al., 2013), making class A an inclusive warning category that also captures borderline interactions. Forty of the 98 class-A alignments reach or exceed the relative hybridization energy of the weakest recovered intended target of the same guide.

These annotations make unavoidable residual predicted risk visible and actionable. They are not intended as numerical probabilities of off-target silencing: guide processing, target accessibility, tissue-specific expression, and other biological variables also influence efficacy. Nevertheless, the underlying WMD determinants are empirically supported, and the ranking provides a practical way to identify the most plausible alternative targets when orthogonal validation of a screen hit fails. Candidate phenotypes should therefore be interpreted together with the construct-specific target list and independent genetic or protein-level validation (Supplementary Fig. S3).

### Next-generation sequencing (NGS) quality control of *pamiR* binary vector library

The physical library was assembled from 2,285 synthesized target-site oligonucleotides, followed by insertion of the MIR319a stem-loop and pooled quality control (Supplementary Fig. S2A). Stem-loop integration was nearly complete (Supplementary Fig. S2B). Next-generation sequencing (NGS) recovered 2,218 of 2,285 designed constructs (97.1%) in the final plasmid pool. Among detected constructs, the mean and median read counts were 65.4 and 62, respectively. The most abundant construct, *pamiR-221*, was represented by 462 reads, approximately sevenfold the mean. In total, 1,882 of 2,218 detected constructs (84.9%) were represented within twofold of the median, indicating a concentrated abundance distribution (Fig. 1G). Read counts for all individual *pamiRs* are provided in Supplementary Dataset S1.

Apparent library coverage is inherently dependent on sequencing depth, as deeper sequencing increases the probability of detecting rare constructs. However, such rare sequences may contribute little to the effective diversity of a subsequent genetic screen if their abundance is orders of magnitude below that of the average construct. In our dataset, the least abundant detected constructs were approximately 60-fold less abundant than the mean, rather than extending into an extremely low-abundance tail. Thus, the 97.1% molecular coverage reflects a library in which nearly all detected constructs remain represented within a comparatively constrained abundance range, increasing their practical likelihood of being sampled during plant transformation and screening. Of note, the NGS quality control only reports on if and to what relative amount a library DNA construct was designed i.e., the relative construct distribution within a plant mutant population obtained by floral-dip is likely different.

The final binary vector which contains the *pamiR* library combines an exchangeable UBQ10 promoter with a HSP terminator, both to achieve constitutive and strong gene expression, and the fluorescence-accumulating seed technology (FAST) marker (Grefen et al., 2010; Nagaya et al., 2010; Shimada et al., 2010; Pratt et al., 2020; Schwartz et al., 2025). UBQ10 was also recently used for constitutive amiRNA expression in the mTACT resource, which discussed potential loss of silencing over generations from 35S-driven constructs commonly driving amiRNA expression (Anfang et al., 2025). FAST allows transformed seeds to be selected by fluorescence before germination, avoiding antibiotic or herbicide treatment prior to the screening (Fig. 1B). This enables direct phenotyping of primary transformants in screens performed with the final vector. Candidate guides are recovered from plants by standardized PCR and Sanger sequencing. To facilitate reproducible use of the pooled resource, the library is distributed together with a detailed handling and amplification guide (Supplementary File S1) and an example vector file (Supplementary File S2).

### Two complementary proof-of-concept screens recover phenotypes expected from redundant plastid gene families

We tested the *pamiR* library in two screening formats. Screen 1 used automated growth and chlorophyll-fluorescence phenotyping to identify altered photosynthetic performance (Figs. 2A, B and S4). Screen 2 used germination in the presence of paclobutrazol (PBZ), an inhibitor of gibberellin (GA) biosynthesis (Hedden and Graebe, 1985), to enrich for lines with altered abscisic acid (ABA)/GA-dependent germination behavior (Figs. 2C and 5A). These screens were intended as proof of principle: the primary output is a candidate *pamiR* line whose guide and phenotype must then be validated independently.

**Figure 2.**
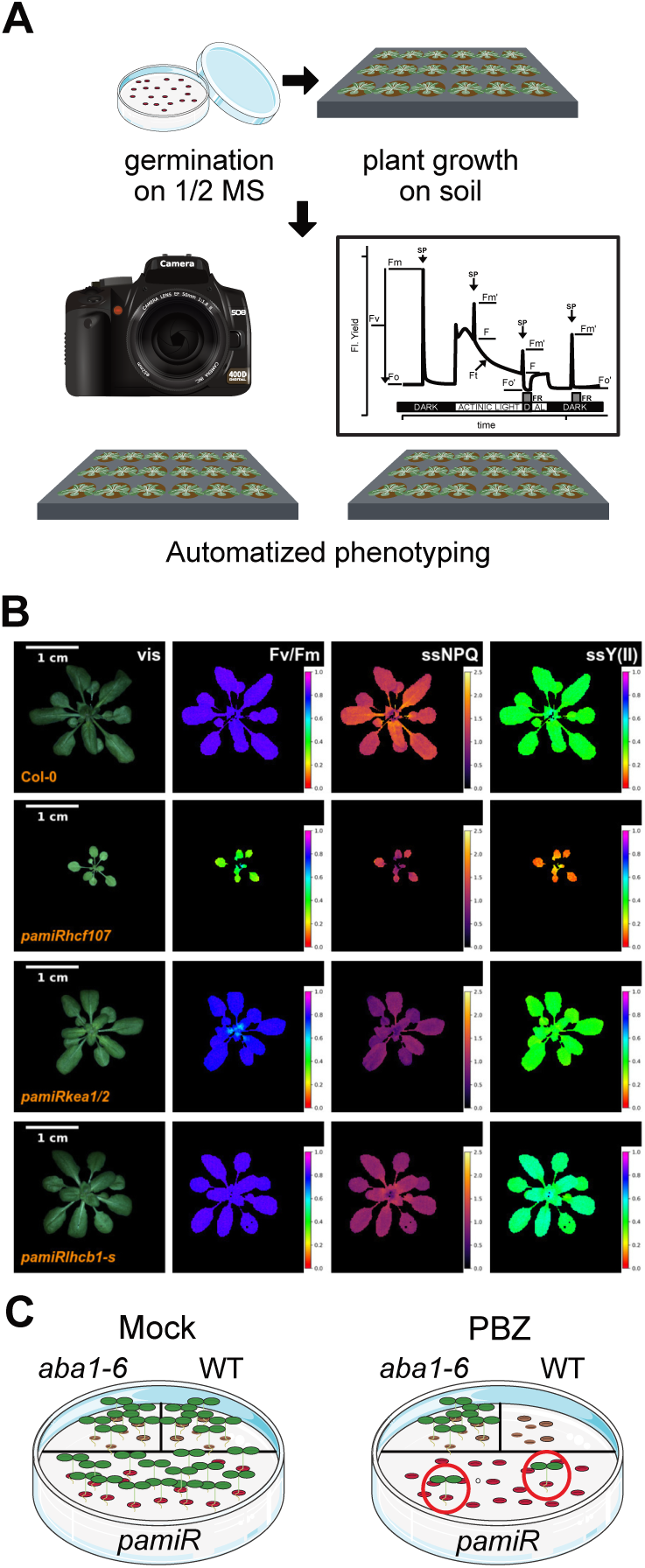
Proof-of-concept *pamiR* screening strategies. A) Workflow for screen 1. Seedlings were established on half-strength MS medium, transferred to soil, and phenotyped by automated RGB imaging and chlorophyll-fluorescence measurements. B) Representative T_2_ lines from screen 1. Columns show visible images, *F*_v_/*F*_m_, steady-state NPQ, and steady-state Y(II), recorded at 185 PAR. WT is Col-0. “-s” denotes a guide designed to target more than two loci. C) Schematic of screen 2, in which WT, *aba1-6*, and *pamiR* seeds were germinated on mock or 40 µM paclobutrazol (PBZ). PBZ inhibits gibberellin biosynthesis and thereby shifts the ABA/GA-dependent germination balance. PBZ-tolerant *pamiR* seedlings were selected as candidates for subsequent characterization.

In screen 1, approximately 600 independent T1 *pamiR* lines were initially screened for altered chlorophyll fluorescence and visible phenotypes using PAM imaging. From these, 100 candidate lines were selected, of which 12 were subsequently subjected to detailed automated phenotyping. The screen recovered multiple lines with phenotypes consistent with previously characterized plastid perturbations (Supplementary Table S1; Supplementary Fig. S4). Three examples are highlighted below. “-s” denotes more than two silenced loci i.e., individuals are expected to resemble triple and higher-order loss-of-function mutants. The ability to recover higher-order-like phenotypes from a single introduced guide is consistent with the intended purpose of multi-target amiRNA design, but phenotypic similarity alone does not prove which family members are depleted.

### *pamiRhcf107* - a viable line affecting an essential PSII-biogenesis factor

*pamiRhcf107* (*pamiR-1990*; Supplementary Table S1) targets HIGH CHLOROPHYLL FLUORESCENT 107 (HCF107; *AT3G17040*), a maturation factor required for accumulation of PsbH-containing PSII complexes (Felder et al., 2001). The line shows chlorosis, slower growth, and reduced *F*_v_/*F*_m_, phenotypes consistent with impaired PSII function (Figs. 2B and S4B-H). Unlike severe *hcf107* loss-of-function alleles, the *pamiR* line remains viable and transmits a stable phenotype. This is consistent with a partial-loss-of-function interpretation, but the extent of HCF107 depletion in this line is not yet established since we could not obtain an HCF107 antibody and should not be inferred from viability alone.

### *pamiRkea1/2* - independent guides converge on KEA1/KEA2

The novel *pamiRkea1/2* allele (*pamiR-401*; Supplementary Table S1) showcases the library’s power to bypass FGR since individual single mutants in either *KEA1* (*AT1G01790*) or *KEA2* (*AT4G00630*) behave like wild-type plants. Targeted knockdown of these specific chloroplast members is necessary because silencing the entire KEA family, which includes proteins localized to the endomembrane system, would likely result in pleiotropic phenotypes. *pamiRkea1/2* mutants recapitulated the expected *kea1kea2* T-DNA double mutant and the previously isolated independent *amiRkea1/2^Kunz2014^*allele phenotypes i.e., diminished growth, low *F*_v_/*F*_m_, and leaf virescence over several generations (Fig. 3A-C; (Kunz et al., 2014)).

**Figure 3.**
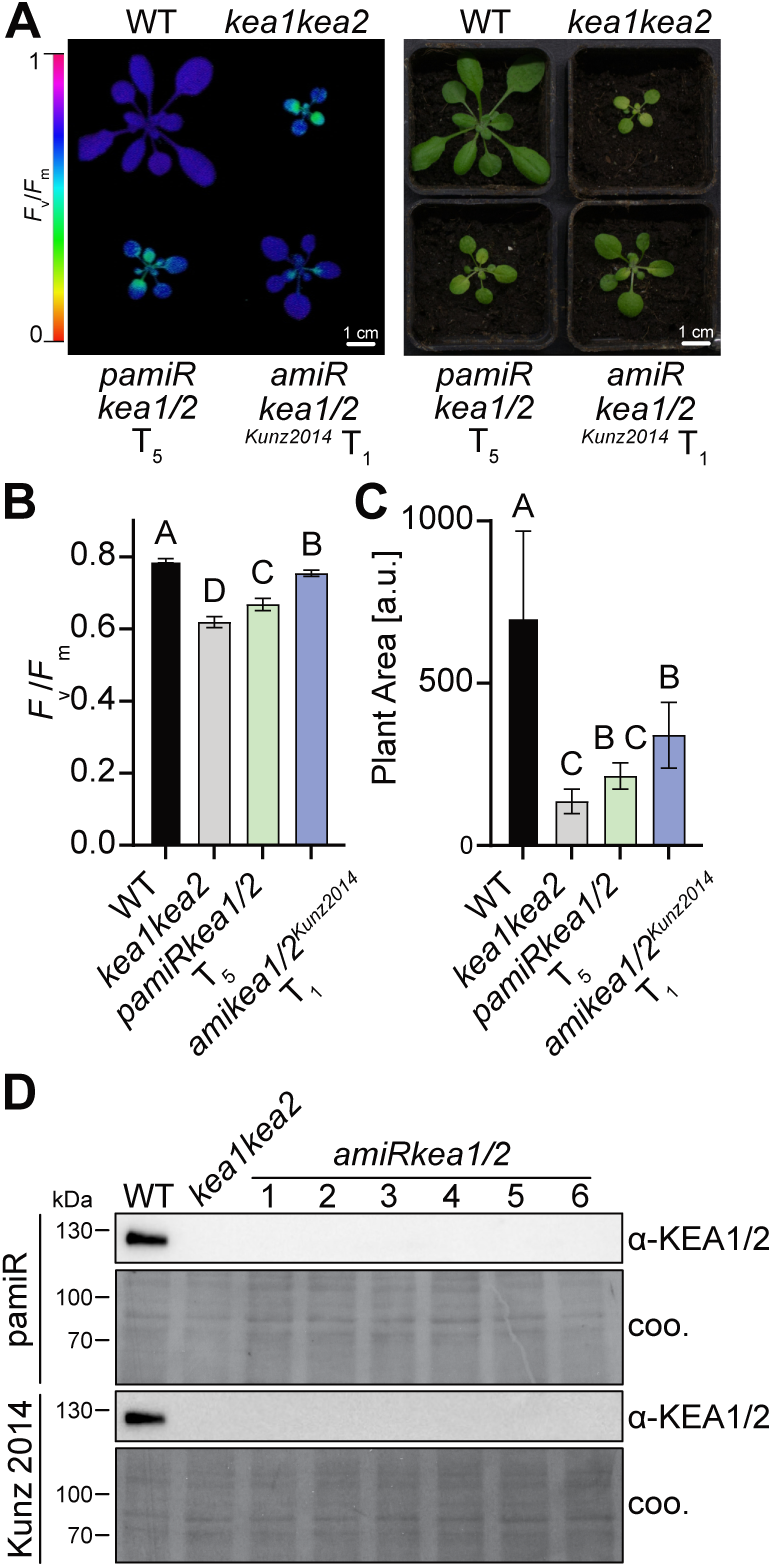
Independent genetic and protein-level validation of *pamiRkea1/2*. A) *F*_v_/*F*_m_ false color and visible images of WT, the loss-of-function *kea1kea2*, *pamiRkea1/2* (T_5_; *pamiR-401*), and the independent *amiRkea1/2^Kunz2014^*line (T_1_). B) Quantification of Fv/Fm. Show are mean values +/− SD (n > 8). Different letters denote statistically distinguishable groups assessed by ordinary one-way ANOVA with Tukey’s multiple comparisons test. C) Quantification of plant area. Show are mean values +/− SD (n > 8). Different letters denote statistically distinguishable groups assessed by ordinary one-way ANOVA with Tukey’s multiple comparisons test. D) KEA1 immunoblots comparing individual plants of *pamiRkea1/2* T_5_ and *amiRkea1/2^Kunz2014^* T_1_ with *pamiR*with WT and *kea1kea2* controls; Coomassie stainings of the same membranes (coo) are shown as a loading controls.

Three observations support attribution of this phenotype to the specific loss of KEA1/KEA2. First, using an KEA1 N-terminal antibody detecting both KEA1 and KEA2 (Bölter et al., 2020), the KEA1/2 immunoreactive signal is below the assay detection limit in the novel *pamiRkea1/2* isolated in this study and in the independent *amiRkea1/2^Kunz2014^* line, as well as in the *kea1-1kea2-1* loss-of-function control (*kea1kea2*), while signal is detected in WT (Fig. 3D). Second, *pamiRkea1/2* (*pamiR-401*) and *amiRkea1/2^Kunz2014^* guide differ at 11 of 21 positions and target different sites in KEA1/KEA2. At the WMD3 design settings used here, both guides return *KEA1* and *KEA2* but no additional locus transcript(s). Third, under a deliberately permissive WMD3 search (target filter disabled, energy cutoff 10%, up to five mismatches), *pamiRkea1/2* returned 46 non-*KEA* genes and the independent guide *amiRkea1/2^Kunz^* 39, with no non-*KEA* transcript common to both returned sets. This does not exclude every conceivable off-target or insertional effect, but it makes a shared predicted direct off-target under the tested search model a highly unlikely explanation for the convergent phenotype.

Importantly, all *amiRkea1/2* mutant lines showed quantitatively weaker defects than the T-DNA allele *kea1kea2*: *F*_v_/*F*_m_ and plant area differ among the lines (Fig. 3B and C) despite KEA1/2 signals being below the immunoblot detection limit. Different residual, spatial, or temporal silencing is a plausible explanation and should be considered when interpreting amiRNA phenotypes. Additionally, insertion/background effects cannot be excluded solely from the immunoblot.

### *pamiRlhcb1-s* - paralogue-resolved proteomics supports target-restricted LHCB1 depletion

*pamiRlhcb1-s* carries *pamiR-983* (Supplementary Table S1), designed to target the following four *LHCB1* loci encoding for the peripheral PSII antenna: *LHCB1.1* (*AT1G29920*), *LHCB1.2* (*AT1G29910*), *LHCB1.3* (*AT1G29930*), and *LHCB1.5* (*AT2G34420*). Fig. 4A summarizes predicted guide-family relationships and is a design schematic rather than an expression measurement. The *pamiRlhcb1-s* line displays chlorosis and reduced regulated non-photochemical energy dissipation, Y(NPQ) (Figs. 2B and S4F). These phenotypes are consistent with previous genetic perturbations of LHCB1: plants with an amiRNA targeting *LHCB1* genes and CRISPR/Cas9-mediated depletion reduced chlorophyll accumulation, pale green phenotypes and reduced NPQ (Pietrzykowska et al., 2014; Sattari Vayghan et al., 2022).

**Figure 4.**
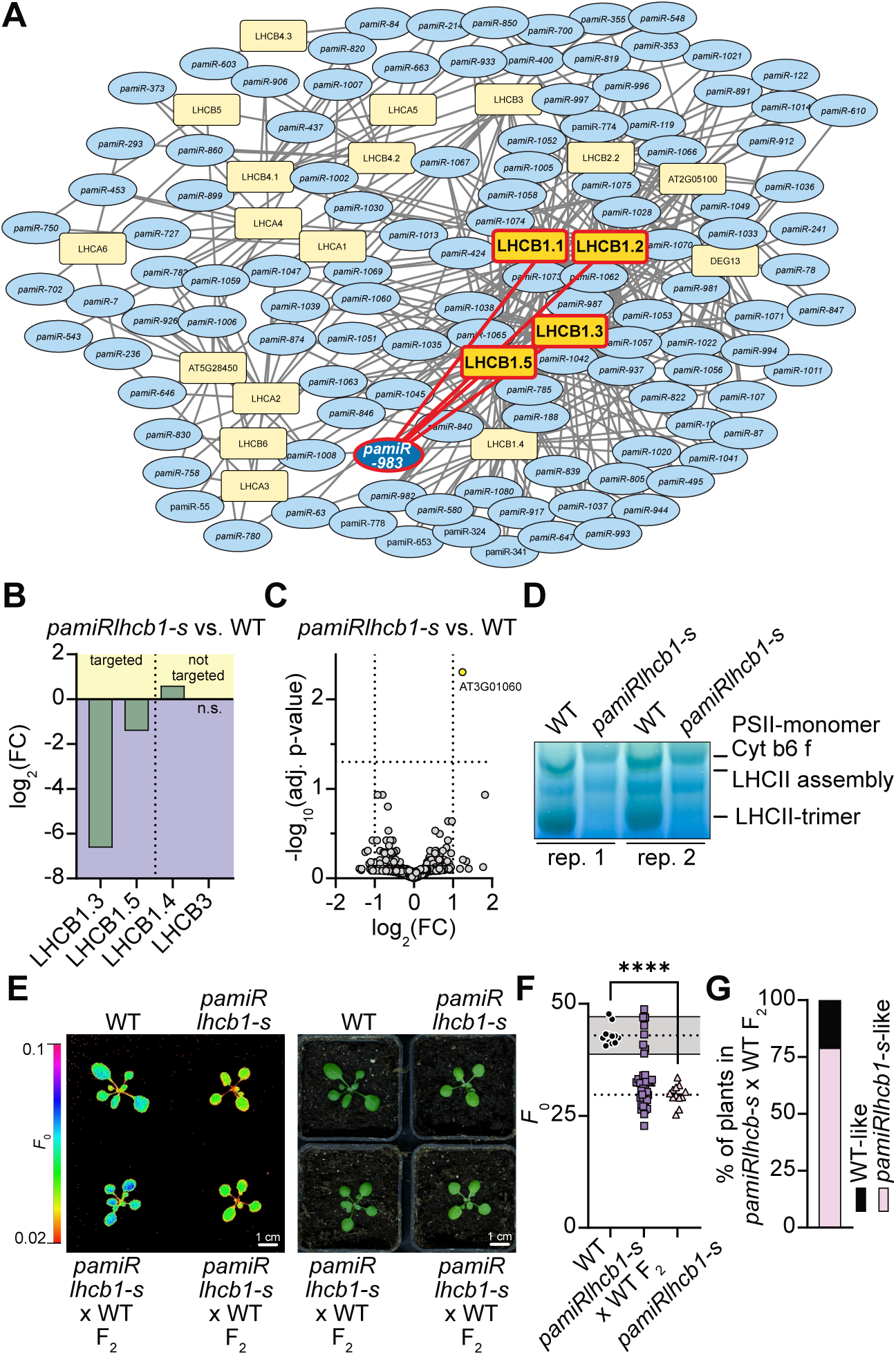
Protein-level, complex-level, and genetic validation of *pamiRlhcb1-s*. A) Designed *pamiR*-LHCB relationships: *pamiR-983* and its four intended targets (LHCB1.1, LHCB1.2, LHCB1.3, and LHCB1.5) are highlighted. B) Paralogue-resolved LHCB abundance from DIA proteomics. *pamiR* targeted LHCB1.3 and LHCB1.5 as well as untargeted LHCB1.4 and LHCB3 were uniquely quantifiable. Significance was assessed by an unpaired t test with Welch correction (n = 4; n.s. = not significant). C) Proteome-wide differential-abundance analysis of *pamiRlhcb1-s* versus WT. Significance was defined as Benjamini-Hochberg-adjusted *P* < 0.05 and |log_2_ fold change| > 1 (n=4) and indicated by dashed lines. D) BN-PAGE of thylakoid complexes from biological replicates 1 and 2; replicates 3 and 4 are shown in Supplementary Fig. S5A. Loading was adjusted to approximate equal fresh-weight equivalents based on chlorophyll content per fresh weight. Regions corresponding to LHCII trimers, higher-order LHCII assemblies, cytochrome *b*6*f*, and PSII monomers are indicated. E) Representative F_0_ false color and visible images of WT, the stable *pamiRlhcb1-s* line, and segregating *pamiRlhcb1-s* × WT F_2_ plants. F) *F*_0_ values of WT, the F_2_ population, and the stable *pamiRlhcb1-s* line. The upper dashed line indicates the WT mean, the gray shaded area two standard deviations around the WT mean, and the lower dashed line the *pamiRlhcb1-s* mean. F_2_ plants with *F*_0_ below the WT mean minus two standard deviations were predefined as *pamiRlhcb1-s*-like. WT and *pamiRlhcb1-s*, n = 12. Significance was assessed by an unpaired t test with Welch correction; ****P < 0.0001. G) Using this threshold, 30/38 F2 plants (79%) were classified as *pamiRlhcb1-s*-like and 8/38 (21%) as WT-like.

To directly assess depletion of the intended targets, while resolving individual paralogues and simultaneously testing for broader protein-level effects, we quantified the *pamiRlhcb1-s* proteome by high sensitivity DIA mass spectrometry with label-free quantification. Because the LHCB1 paralogues are highly similar, conventional protein-group-level quantification cannot reliably distinguish them: most of the detected peptides are shared between isoforms, and the resulting protein-group abundances therefore combine information from several LHCB1 proteins. Since our aim was to determine whether the individual LHCB1 proteins targeted by *pamiRlhcb1-s* were depleted, we instead performed the LHCB analysis at the precursor level and restricted isoform-specific quantification to peptides that could be assigned unambiguously to a single LHCB family member, as detailed in Materials and Methods (Supplementary Table S2). Among the four intended targets, LHCB1.3 (AT1G29930) and LHCB1.5 (AT2G34420) were uniquely quantifiable and were depleted (log_2_ fold changes −6.6 and −1.4, respectively; Fig. 4B). LHCB1.1 and LHCB1.2 could not be resolved by unique peptides. In the same samples, the quantifiable untargeted family members LHCB1.4 (AT2G34430) and LHCB3 (AT5G54270) were not significantly changed. Thus, the experiment provides a paralogue-resolved internal specificity test within the LHCB family (Fig. 4B; Supplementary Table S2).

The same DIA dataset also provides a proteome-wide test for unintended protein depletion in the assayed tissue and condition. After filtering, 9,441 non-LHCB protein groups were included in the genome-wide differential-abundance analysis (Supplementary Dataset S3). No significantly decreased protein was detected, whereas AT3G01060 (lysine-tRNA ligase) was the only significantly increased protein (Fig. 4C). AT3G01060 is not returned as a *pamiRlhcb1-s* target even in the deliberately permissive WMD3 query used here. In addition, a transcript increase is unexpected for direct amiRNA-mediated silencing. Together with the unchanged untargeted LHCB paralogues, this provides evidence against widespread strong protein-level off-target silencing in this line, while not excluding effects on undetected proteins or smaller changes below the statistical threshold (Fig. 4C).

We next asked whether the protein depletion detected by DIA was accompanied by the characteristic changes in LHCII organization previously associated with LHCB1 deficiency. BN-PAGE revealed reduced accumulation of LHCII trimers and higher-order LHCII assemblies across four biological replicates (Figs. 4D and S5A). This closely resembles previously reported results (Pietrzykowska et al., 2014). Together with the chlorosis and reduced Y(NPQ), these orthogonal phenotypes corroborate the LHCB1 depletion measured by DIA proteomics.

In addition to these previously established consequences of LHCB1 depletion, *pamiRlhcb1-s* showed a pronounced reduction in *F*_0_ (minimum fluorescence yield; Fig. 4E and F). We took advantage of the strong separation in *F*_0_ between *pamiRlhcb1-s* and WT as a robust quantitative marker for the subsequent segregation analysis. To test whether a representative *pamiR*-induced phenotype behaves dominantly, a stable *pamiRlhcb1*-s line was backcrossed to WT. PCR-positive F_1_ progeny were selfed and 38 next generation F_2_ plants were phenotyped together with WT and stable *pamiRlhcb1-s* controls. F_2_ plants with *F*_0_ below the WT mean minus two standard deviations were classified as *pamiRlhcb1-s*-like. Using this predefined threshold, 30 of 38 F_2_ plants (79%) were classified as *pamiRlhcb1-s*-like and 8 of 38 (21%) as WT-like (Fig. 4E-G). The observed segregation is consistent with a dominant rather than recessive phenotype.

### A PBZ germination screen identifies *pamiRnced-s*, a guide designed to co-target five *NCED* genes

In screen 2, approximately 600 independent *pamiR* T_1_ lines were germinated on half-strength MS medium containing 40 µM PBZ. PBZ inhibits gibberellin biosynthesis; reduced GA promotes dormancy. Hence, ABA-deficient mutant seeds show increased germination under PBZ treatment (Barrero et al., 2005; Lefebvre et al., 2006). The ABA synthesis mutant *aba1-6* served as positive control. We recovered a *pamiR* line with PBZ-tolerant germination that remained detectable in subsequent generations (Figs. 2C and 5A). Guide identification showed that this line carries *pamiR-1034* (Supplementary Table S1), designed to target 9-cis epoxycarotenoid dioxygenase (NCED) loci: *NCED2* (*AT4G18350*), *NCED3* (*AT3G14440*), *NCED5* (*AT1G30100*), *NCED6* (*AT3G24220*), and *NCED9* (*AT1G78390*; Fig. 5B and Supplementary Table S3). NCED enzymes cleave violaxanthin or neoxanthin to yield the ABA precursor Xanthoxin. Interestingly, *nced6* and *nced9* single mutant seedings show only moderate PBZ tolerance whereas only *nced6nced9* double mutants are PBZ resistant (Lefebvre et al., 2006). Therefore, our results on the *pamiRnced-s* allele emphasize the power of the *pamiRs* to overcome FGR.

**Figure 5.**
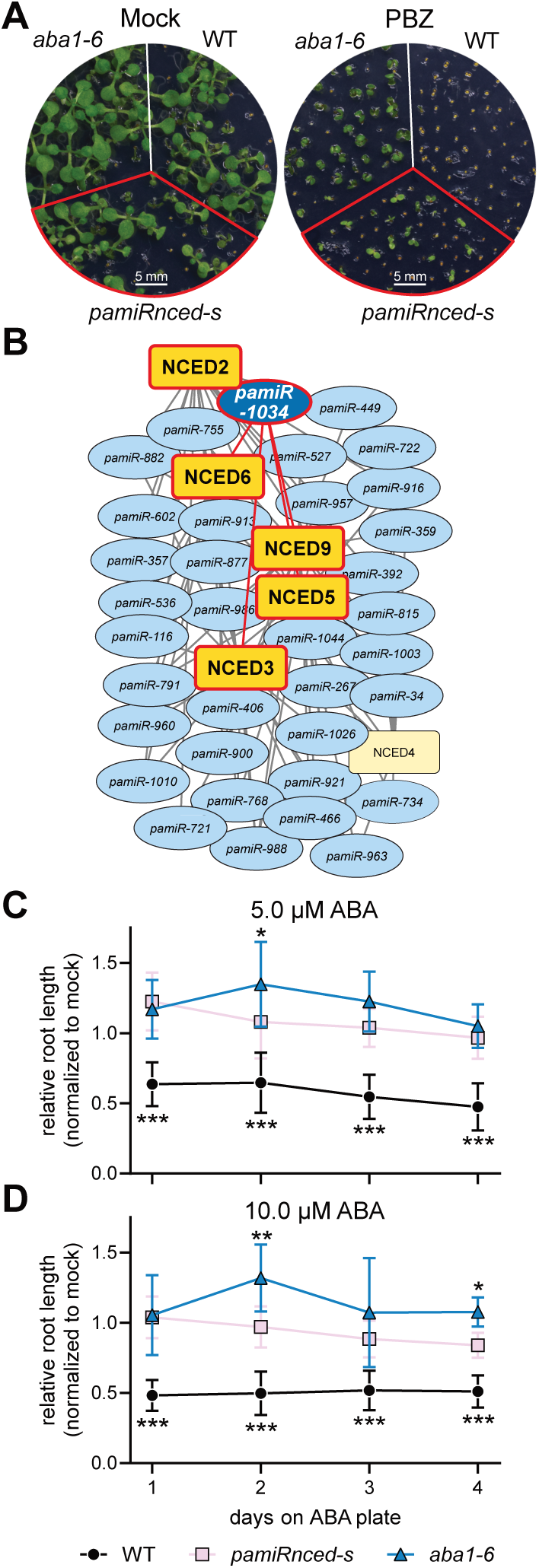
Target assignment and ABA-related physiological characterization of *pamiRnced-s*. A) Representative mock and 40 µM PBZ plates containing WT, *aba1-6*, and *pamiRnced-s*. The plate phenotype is presented as qualitative screening evidence. B) Designed *pamiR*-*NCED* relationships; *pamiR-1034* and its five intended targets *NCED2*, *NCED3*, *NCED5*, *NCED6*, and *NCED9* are highlighted. The network represents designed guide-target relationships rather than measured transcript or protein depletion. C-D) Relative root elongation after transfer to 5 µM ABA (C) or 10 µM ABA (D), normalized to the corresponding mock-grown controls. Data are shown as mean +/− SD for n ≥ 8 biological replicates per genotype. Statistical differences were assessed by two-way ANOVA with Sidak multiple-comparisons correction. Asterisks plotted below the WT series indicate significant differences between WT and both *pamiRnced-s*/*aba1-6* at the indicated time point; asterisks above the graphs indicate significant differences between *pamiRnced-s* and *aba1-6*. *P < 0.05, **P < 0.001, ***P < 0.0001.

The five-target assignment is computational and can be inspected directly from the guide sequence. In Supplementary Table S3, the five reported targets contain two to four mismatches to *pamiR-1034*, no mismatch at guide positions 10/11, and WMD3 relative hybridization energies of 73.04-98.26%; all five are returned at the stated design settings. The four other NCED family members contain seven to nine mismatches and are not returned at those settings; three also contain a mismatch at guide position 10 or 11. We therefore describe these five genes as the predicted intended targets of *pamiR-1034* rather than as experimentally demonstrated depleted proteins.

Direct protein-level confirmation is currently unavailable for this line: the relevant NCED proteins were not detected in the analyzed leaf DIA dataset (Supplementary Dataset S3), and no suitable isoform-resolved antibodies are available. We therefore assessed ABA responsiveness with a root-elongation assay after transfer to 5 or 10 µM exogenous ABA, a strong root growth inhibitor. Relative root growth was normalized to mock treatment. *pamiRnced*-s and *aba1-6* differed from WT i.e., were less ABA sensitive, at multiple time points, while smaller differences between *pamiRnced*-s and *aba1-6* were also detected at selected time points (Figs. 5C and D and S5B-D).

The original PBZ germination phenotype is treated qualitatively. Preliminary control experiments showed that discrimination between WT and *aba1-6* occurs within a narrow PBZ concentration window and is sensitive to seed age and batch. We therefore calibrate the screening concentration with age-matched WT and *aba1-6* controls before each screen/assay rather than interpreting a broad PBZ concentration series as a quantitative phenotype. The plate image in Fig. 5A is shown as proof-of-concept screening evidence, whereas the root-elongation experiment (Fig. 5C and D) provides the quantitative physiological characterization.

## Conclusions and limitations

*pamiR* is a powerful organelle-specific screening resource rather than a substitute for definitive causal genetics. The full library has been computationally reassessed construct by construct, and representative proof-of-concept plant mutant lines have been tested experimentally at protein, complex, physiological, and genetic levels, including detailed analyses of LHCB1 and KEA1/KEA2. The characteristic *pamiRlhcb1*-s phenotype remained detectable across generations and after backcrossing and selfing. At the same time, the KEA comparison illustrates an important limitation: the two amiRNA lines and the *kea1kea2* T-DNA loss-of-function line differ quantitatively in *F*_v_/*F*_m_ and plant area even though KEA1 immunoreactive signal is below the detection limit in all three perturbed genotypes. Different residual or spatiotemporal silencing is a likely explanation, but insertion/background effects and other biological differences must also be considered. As shown in the example of *HCF107*, where a full loss-function allele does not produce seeds, the milder defects in *pamiR* lines that yield homozygous seed pools for in-depth follow-up studies can be a desirable outcome.

Every *pamiR* primary screen hit should be validated by multiple, non-mutually-exclusive approaches. Useful steps include testing persistence of the phenotype in descendants of the same transformation event; backcrossing a primary *pamiR* line to the WT background and reassessing the phenotype in segregating descendants to reduce transformation/insertion-background effects; testing a second sequence-independent amiRNA against the same target set, readily designed and screened in WMD3, to lower the probability of a shared sequence-dependent off-target; comparing independent null alleles such as CRISPR or T-DNA lines where genetic redundancy permits; complementation or rescue; and direct verification of target-protein depletion where technically feasible. If an orthogonal strategy fails to reproduce a phenotype, the construct-specific WMD3 alignment list provides a defined set of alternative predicted targets to investigate.

The *pamiR* resource is publicly available as DNA stocks. It is accompanied with a detailed handling guide that describes pooled-library amplification, *Agrobacterium* transformation, plant screening, guide recovery, and validation (Supplementary File S1), together with the per-construct sequence/read-count/off-target annotation (Supplementary Dataset S1) and an example vector file (Supplementary File S2). For practical screens we recommend ≥10,000 independent primary transformants, corresponding to approximately 4.4-fold nominal representation of the 2,285-guide library, to approach broad coverage. However, substantially smaller, non-saturating screens can still yield informative candidates: the proof-of-concept screens performed here at a scale of approximately 600 T_1_ individuals recovered multiple interpretable phenotypes, including *pamiRkea1/2*, *pamiRlhcb1-s*, and *pamiRnced-s*. Thus, the 10,000-transformant recommendation should not be interpreted as a minimum screen size, literal saturation, or a guarantee that every guide has been sampled. Random sampling and the measured non-uniform construct abundance mean that some *pamiRs* can remain unseen even at 10,000 transformants. Actual screen saturation therefore has to be judged empirically from recovered-guide frequencies and phenotype discovery rather than from transformant count alone.

Finally, the library-wide target assignments remain computational predictions for untested constructs, WMD3 does not capture all biological determinants of silencing, and the plastid target set inherits limitations of the localization datasets used for its construction. The experimental examples validate the utility of the resource, not every individual *pamiR*. These limitations are why *pamiR* is intended to narrow and accelerate plastid-focused forward-genetic discovery while retaining independent validation as an essential final step.

In summary, our study shows that the application of FGR-free organelle-specific forward genetics is enabled through the design of multi-target amiRNAs joint together in subcellular libraries. The ever-improving sensitivity of mass-spectrometry and orthogonal high resolution fluorescence protein localization studies will likely complete the cellular protein map for *A. thaliana* soon i.e., additional organelle-specific amiRNAs can be designed following our straight-forward approach. An obvious next candidate is the mitochondrion, for which detailed proteome information is now available (Fuchs et al., 2020; Rugen et al., 2024). The described library here is accessible to all interested via the community stock centers. We hope *pamiR* will enable many researchers to unveil new gene functions further by closing the gap of a function genome understanding for the model plant Arabidopsis.

## Materials and Methods

### Plant material and growth conditions

*Arabidopsis thaliana* Columbia-0 (Col-0; hereafter WT) was used as the wild type, and all lines were in the Columbia background unless otherwise stated. The ABA-deficient *aba1-6* mutant (Koornneef et al., 1982; Niyogi et al., 1998) served as a positive control in the PBZ and ABA-response experiments. The KEA loss-of-function plant line *kea1-1kea2-1* (in short *kea1kea2*) and *amiRkea1/2^Kunz2014^* were described by (Kunz et al., 2014). Unless otherwise specified, seeds were surface sterilized with 70% ethanol, plated on half-strength Murashige and Skoog (½ MS) medium ((Murashige and Skoog, 1962); Duchefa Biochemie) containing 0.8% phytoagar, and stratified for at least 2 d at 4°C in darkness. Seedlings were grown for 7-9 d under 16 h light at 22°C and 100 µmol photons m⁻² s⁻¹ with 30% relative humidity / 8 h dark at 18°C and 30% relative humidity, transferred to soil, and grown for a further 14-16 d under 16 h light at 22°C and 100 µmol photons m⁻² s⁻¹ with 55% relative humidity / 8 h dark at 18°C and 65% relative humidity. Screen 1 used the separate growth regime described below.

### In silico design of the plastid amiRNA (*pamiR*) library

To assemble a plastid-enriched target list, we surveyed six published proteomic datasets from chloroplasts or plastid membranes (Sun et al., 2008; Ferro et al., 2010; Simm et al., 2013; Tomizioli et al., 2014; Lundquist et al., 2017; Trentmann et al., 2020). Candidates were assigned a provisional support score according to their recurrence across these datasets, and the resulting collection was subsequently refined using additional localization resources and literature, including SUBA and its Multiple Marker Abundance Profiling (MMAP) tool (Schwacke et al., 2003; Hooper et al., 2014; Hooper et al., 2017).

The resulting nuclear gene list was compared with the PHANTOM amiRNA resource to identify existing family-targeting designs (Hauser et al., 2013). Designs that also targeted non-plastid family members according to the selected annotations were excluded or redesigned with the WMD strategy (Schwab et al., 2006; Ossowski et al., 2008). Of 2,281 distinct amiR sequences in the final collection, 679 are shared with PHANTOM; 672 of these have the same annotated target-gene set in both collections. The remaining sequences were selected/designed to complete coverage of the curated plastid target set. Supplementary Fig. S1B visualizes guide-gene relationships but should be interpreted as a design network, not measured silencing.

### amiRNA design rules

Artificial microRNAs were designed with the Web MicroRNA Designer (WMD) framework used for PHANTOM (Schwab et al., 2006; Ossowski et al., 2008). Candidate 21-nt guides were selected using WMD sequence/duplex criteria, including guide-strand asymmetry, stringent complementarity in the 5-prime/central region, no intended-target mismatch at positions 10/11 under the standard design rule, and predicted duplex-energy criteria. These rules were derived from experimental analyses of plant miRNA/amiRNA targeting and were subsequently supported by protein-based amiRNA screens (Schwab et al., 2006; Li et al., 2013). For intended family targets, WMD’s multi-target strategy was used so that one guide had to satisfy the target-recognition criteria for every member assigned to that guide.

Gene-family groupings were therefore operational rather than based on a fixed global sequence-identity cutoff: paralogues were co-targeted when one 21-nt guide could satisfy the design criteria for all of them, and families were divided into smaller co-targetable subsets when necessary. The resulting collection contains 2,285 amiRNAs (2,281 unique guide sequences) directed against 2,202 unique intended genes, with 1.72 intended genes per amiRNA on average (median 1; range 1-9). Corresponding amiR/amiR* sequences were inserted into the Arabidopsis MIR319a/pRS300 architecture in a pGreen-based vector (Schwab et al., 2006; Schwenkert et al., 2023).

### Off-target prediction and assignment of specificity classes

To predict off targets based on the most recent Arabidopsis thaliana genome annotation, all 2,285 current-library guide sequences were re-queried with WMD3 Target Search against Araport11 cDNA reference implemented by the platform (release 201606). The query used up to five mismatches, the miRNA target filter, a minimum relative hybridization-energy threshold of 70% of the perfect-match value, folding temperature 23°C, RNAcofold, no gaps, reverse-strand matching, and isoforms collapsed to the best-scoring alignment per gene. Recovered genes were classified as declared intended targets or additional predicted transcript matches (Supplementary Dataset S1) Mismatch positions were indexed from the guide 5-prime end. The complete 4,862 recovered alignments (3,834 intended-target alignments and 1,028 additional predicted alignments), including mismatch positions and relative hybridization energies, are provided in Supplementary Dataset S2.

For the *pamiR* specificity classification, each additional WMD3-predicted alignment was assigned to one of five ordered risk classes using a strict hierarchical cascade of sequence and duplex features known to influence plant miRNA/amiRNA silencing efficiency (Supplementary Table S4). The classification proceeds by testing the strongest mechanistic constraints first: an alignment containing a mismatch at guide position 10 or 11 is assigned to class D (negligible predicted risk); if positions 10/11 are matched, alignments containing two or more mismatches within positions 2–12 are assigned to class C (low); among the remaining alignments, those with relative hybridization energy below 75% of the perfect-match duplex are assigned to class B (moderate), whereas alignments reaching ≥75% are assigned to class A (high). Class E denotes *pamiRs* for which no additional transcript was returned under the stated WMD3 search conditions. Thus, the cascade first evaluates target-pairing geometry and only subsequently duplex stability: a strong predicted duplex does not receive a high-risk classification when its mismatch pattern is expected to strongly impair canonical AGO-mediated cleavage. The resulting classes therefore represent decreasing predicted compatibility with efficient amiRNA-mediated silencing in the order A > B > C > D, with E indicating no predicted off-target under the applied search settings. Each *pamiR* was assigned the class of its highest-risk predicted off-target. The 75% boundary between classes A and B was deliberately chosen below the >80% relative hybridization energy associated with optimal amiRNA activity in protein-based validation experiments (Li et al., 2013), making class A an inclusive warning category rather than a quantitative probability of off-target silencing. The criteria, mechanistic rationale, and numbers of off-target alignments and *pamiRs* assigned to each class are summarized in Supplementary Table S4.

### Molecular cloning of the *pamiR* library

DNA oligomers were synthesized using Agilent SurePrint pooled oligo synthesis (Agilent Technologies, Santa Clara, CA, USA) as 169 bp fragments with amiR and amiR* for each *pamiR*. The oligomers did not include the stem-loop of the amiRNA. A pGreen-based amiRNA vector (Schwenkert et al., 2023; Schwartz et al., 2025) and an oligomer library were amplified by PCR using Platinum SuperFi II DNA Polymerase (ThermoFisher Scientific) and primers that enabled seamless Golden Gate cloning via BsaI (Supplementary Table S5). The amplified vector was gel-purified, while the amplified library PCR was directly purified using the NucleoSpin Gel and PCR Clean-up Mini kit (Macherey-Nagel, Düren, Germany). 20 ng of the PCR-amplified library DNA were inserted into 300 ng of the PCR-amplified plasmid (2:1 molar ratio) through a Golden Gate reaction containing 1x T4 ligase buffer (NEB), 1000 units T4 Ligase (NEB), and 30 units BsaI (ThermoFisher Scientific) in a 40 µl reaction supplemented with 5 mM DTT and 0.5 mM ATP. Incubation was carried out in a PCR cycler with the settings: 30 sec at 37°C, 25 cycles at 5 min at 37°C, followed by 5 min at 16°C, 5 min at 50°C, and 5 min at 80°C. The reaction was directly transformed into Stellar chemically competent cells for cloning (Takara). The transformation reactions were plated on LB-Agar supplemented with 50 µg/ml KANAMYCIN, resulting in more than 20,000 individual colonies carrying the bulk plasmid library without stem-loop (*pamiR*-NoSL). Cells were resuspended in liquid LB supplemented with 50 µg/ml KANAMYCIN using a cell scraper, transferred into 500 ml of supplemented media, grown at 37°C for 3.5 h, and plasmid DNA was extracted from 4 ml of cell culture using the NucleoSpin Plasmid Mini kit (MACHEREY-NAGEL).

Next up, the amiRNA stem-loop was inserted into the bulk plasmid library *pamiR*-NoSL by a second Golden Gate reaction: 200 ng of *pamiR*-NoSL, 200 ng of stem-loop vector (2:1 molar ratio), 1x T4 ligase buffer (NEB), 500 units T4 Ligase (NEB), and 30 units BsaI (ThermoFisher Scientific) in a 20 µl reaction supplemented with 5 mM DTT and 0.5 mM ATP. The Golden Gate reaction took place in a PCR cycler with the following program: 30 s at 37°C, 25 cycles at 5 min at 37°C, followed by 5 min at 16°C, 60 min at 37°C after the addition of 10 units of NotI (NEB), 5 min at 50°C, and 5 min at 80°C. The reaction mix was directly transformed into Stellar chemically competent cells for cloning (Takara). Transformation reactions were plated on LB-Agar supplemented with 50 µg/ml KANAMYCIN, resulting in more than 30,000 individual colonies. Cells were resuspended in liquid LB supplemented with 50 µg/ml KANAMYCIN using a cell scraper, transferred into 250 ml of supplemented media, grown at 37°C for 2 h, and plasmid DNA was extracted from 4 ml of cell culture using the NucleoSpin Plasmid Mini kit (MACHEREY-NAGEL) four times.

Sequencing revealed that the starting amiRNA vector was over-represented in this plasmid pool. The initial amiRNA vector contained two HindIII sites with a 1000 bp distance, while the *pamiR* vectors only contained one HindIII site. This was used to greatly reduce the starting vector contamination: 9 µg of the vector pool was digested with HindIII and ran on a 0.5% (w/v) Agarose gel for 45 min separating the *pamiR* and the starting vector. The single-cut *pamiR* bands were excised and gel-purified using the NucleoSpin Gel and PCR Clean-up Mini kit (MACHEREY-NAGEL). 130 ng of the resulting linearized vector was ligated with T4 ligase (NEB) in a 20 µl standard reaction supplemented with 5 mM DTT and 0.5 mM ATP. The ligation was heat-shock transformed into Stellar chemically competent cells for cloning (Takara) and cells were plated on LB-Agar supplemented with 50 µg/ml KANAMYCIN, resulting in more than 30,000 individual colonies. Cells were resuspended in liquid LB supplemented with 50 µg/ml KANAMYCIN using a cell scraper, transferred into 250 ml of supplemented media, grown at 37°C for 2 h, and plasmid DNA was extracted from 4 ml of cell culture using the NucleoSpin Plasmid Mini kit (MACHEREY-NAGEL) eight times.

### Bioanalyzer measurements

To verify complete stem loop insertion 2.5 µg *pamiR* were digested with BamHI and XmaI at 37°C over night. The digestion was purified using the NucleoSpin Gel and PCR Clean-up Mini kit (MACHEREY-NAGEL). The elution was diluted to achieve a concentration of 10 ng/µl. 1 µl was examined using the High Sensitivity DNA Kit on a Bioanalyzer (Agilent).

### Next-generation sequencing (NGS) of bulked *pamiR* plasmid library

The amiRNA-containing regions in the *pamiR* were PCR-amplified (20 cycles) using Platinum SuperFi II DNA Polymerase (ThermoFisher Scientific) and primers (P5_NGS_f and P7_NGS_r; Supplementary Table S5) to add Illumina sequencing adapters. PCR products were purified using the NucleoSpin Gel and PCR Clean-up Mini kit (MACHEREY-NAGEL) and sequenced 330 bp paired-end on a MiSeq Sequencing System (Illumina) at LMU’s Genomics Core Facility.

Reads were processed in CLC Genomics Workbench (version 20.0.4). Because constructs only differ in the amiR/amiR* regions and can be highly similar, stringent mapping was used to reduce ambiguous assignments: the mismatch penalty was set to 5 and stand-alone read mappings were retained. Coverage and construct read-count distributions were calculated from the resulting per-construct table and plotted in GraphPad Prism (v10.6.1). Of 2,285 designed constructs, 2,218 were detected (97.1%). For detected constructs, the maximum, mean and median read counts were 462, 65.4 and 62, respectively; the coefficient of variation was 47.3%, and 1,882 of 2,218 constructs (84.9%) fell within 0.5-to 2-fold of the median. Per-construct read counts are reported in Supplementary Dataset S1.

### Subcellular distribution of proteins

For the contextual compartment counts in Fig. 1A, all 26,882 Araport11 gene identifiers were queried in SUBA and filtered to retain entries with a single SUBAcon consensus assignment, resulting in 25,972 proteins. The compartment counts shown in Fig. 1A are these mutually exclusive single-SUBAcon assignments and sum to 25,972. SUBAcon integrates experimental localization evidence with prediction, and the counts are therefore consensus annotations rather than uniform direct localization measurements.

### Network construction

*pamiR* IDs and their targets were used to construct networks in Cytoscape (v3.10.4).

### Plant transformation

*pamiR* constructs were transformed into A. tumefaciens GV3101 carrying the pSOUP helper plasmid by electroporation. Successfully transfected cells were selected on LB-Agar supplemented with RIFAMPICIN (20 µg/ml), GENTAMYCIN (10 µg/ml) and KANAMYCIN (50 µg/ml). A. tumefaciens cells carrying the *pamiR* library were grown to an OD600 = 1 and resuspended in 5% sucrose with 0.05% Silwet. A. thaliana inflorescences were dipped into the A. tumefaciens cell suspension for a duration of 30 s. The floral-dip plant transformation was repeated two weeks later to increase the number of *pamiR* carrying T1 individuals.

### *pamiR* proof of concept forward genetic screens

Screens were performed on primary transformants. For screen 2 using the final *pamiR*, T_1_ seeds are selected by FAST fluorescence before germination. Screen 1 used an earlier vector backbone carrying the same amiRNA sequences as the current library and primary transformants were selected on half-strength MS medium containing 15 µg ml⁻¹ HYGROMYCIN. PENDING: insert the exact number of independent primary transformants screened in each screen and the numbers carried forward/confirmed, both here and in Results. For future use of the pooled resource, at least 10,000 independent primary transformants are recommended as a practical lower bound for broad library coverage; this is not literal saturation and does not guarantee recovery of every guide.

### Screen Nr. 1: Photosynthesis and growth phenotyping

One-week-old *pamiR* plants were transferred to soil and grown under the screen-1-specific regime of 12 h light (120 µmol photons m⁻² s⁻¹, 22°C) / 12 h dark (18°C). Primary T1 plants were initially screened for altered chlorophyll fluorescence and visible phenotypes using a Walz Imaging-PAM system after dark adaptation. Selected candidate lines were subsequently subjected to detailed phenotyping in the WSU PhenoCenter from two weeks of age, with daily RGB and chlorophyll-fluorescence measurements for nine consecutive days following the established platform protocol (Li et al., 2021). Images were analyzed with the ImagingPAM pipeline (Schneider et al., 2019). For the quantitative comparisons shown in Supplementary Fig. S4B-G.

### Screen Nr. 2: paclobutrazol-based germination screen

Surface-sterilized T_1_ *pamiR*, WT, and *aba1-6* seeds were plated on half-strength MS medium containing 0.8% phytoagar, with or without 40 µM PBZ prepared from a 10 mM stock in ethanol. Plates were stratified for 3 d before transfer to the growth chamber, and candidate PBZ-tolerant seedlings were transferred to soil. Because PBZ-dependent germination is sensitive to seed age/batch and the discriminatory concentration range is narrow, the working PBZ concentration was established in preliminary control plates using age-matched WT and *aba1-6* before the screen/assay. The PBZ plate phenotype is therefore used qualitatively as a screening readout rather than as a quantitative dose-response phenotype.

### Identification of gene(s) targeted by *pamiRs* in screen Nr. 1 and Nr. 2

Candidate guide sequences were recovered from genomic DNA by PCR across the amiR/amiR* region and Sanger sequencing, then matched to the *pamiR* sequence table to identify the construct and its declared target set. The screen protocol used mir319_f (5’-CAAACACACGCTCGGAC-3’) and HSP18_r (5’-CACAAATTCATAACACAACAAGC-3’), yielding a 517-bp product; the handling manual also provides an alternative UBQ10_f/mir319_rev primer pair (Supplementary File S1). Primer sequences are retained in Supplementary Table S5. Identifying the guide carried by a line identifies its designed target set but does not by itself demonstrate depletion of every declared target.

### Quantitative DIA proteomics

Four biological replicates per genotype were analyzed. Frozen rosette leaf tissue was ground and 50 mg powder was resuspended in 100 mM HEPES (pH 7.5), 150 mM NaCl, 10 mM DTT, 6 M guanidine chloride, and cOmplete Protease Inhibitor Cocktail (Roche). Samples were sonicated with a Branson Sonifier B-12, incubated at 60°C for 10 min, and centrifuged at 10,000 x g for 15 min. Proteins were precipitated by chloroform-methanol extraction, washed with methanol, dried, and resuspended in 3 M urea, 2 M thiourea and 50 mM HEPES (pH 7.8). Protein concentration was measured with the Pierce 660 nm Protein Assay. Aliquots of 100 ug protein were reduced with 10 mM DTT for 30 min at 37°C, alkylated with 50 mM iodoacetamide for 20 min at room temperature in the dark, and digested overnight at 37°C with 1 ug trypsin. Peptides were purified on C18 StageTips (Rappsilber et al., 2003), and 200 ng was analyzed by LC-MS/MS.

LC-MS/MS was performed on a nanoElute2 system coupled to a timsTOF HT mass spectrometer (Bruker Daltonics) via a CaptiveSpray nano-ESI source. Peptides were trapped on a PepMap Neo C18 column (300 um x 5 mm, 5 um; Thermo Fisher Scientific) and separated on a PepSep Ultra C18 column (25 cm x 75 um, 1.5 um; Bruker Daltonics) using a 60-min linear 5-37% (v/v) acetonitrile gradient at 250 nl min-1 and 50°C. diaPASEF acquisition used the DIA PASEF-short-gradient method with a 100-1700 m/z mass range, 0.85-1.3 Vs cm-2 ion-mobility range, and 100-ms ramp time.

#### Mass spectrometry data processing

##### Spectral library generation and database search (FragPipe)

The complete acquisition set comprised 12 ddaPASEF and 12 diaPASEF runs. Raw timsTOF data (.d) were processed in FragPipe v23.1 using the DIA_SpecLib_Quant_diaPASEF workflow (Yu et al., 2023). Pseudo-MS/MS spectra were extracted from the diaPASEF files with diaTracer v2.2.1 (Li et al., 2025), and the resulting pseudo-spectra together with the ddaPASEF runs were searched with MSFragger v4.4.1 (Kong et al., 2017; Teo et al., 2021) against Araport11_genes.201606-pep.fasta supplemented with plastid- and mitochondrial-encoded proteins incorporating known RNA edits (van Wijk et al., 2023), common contaminants, and reversed decoy sequences. Searches used tryptic specificity with up to two missed cleavages, precursor and fragment mass tolerances of ±20 ppm, carbamidomethylation of cysteine as a fixed modification, and oxidation of methionine and protein N-terminal acetylation as variable modifications. Peptide-spectrum matches were rescored with MSBooster v1.3.17 (Yang et al., 2023) and Percolator v3.7.1 (Käll et al., 2007) and filtered to 1% PSM- and protein-level FDR with ProteinProphet (Nesvizhskii et al., 2003) in Philosopher v5.1.2 (da Veiga Leprevost et al., 2020). A single empirical spectral library was built from all identified peptides with EasyPQP v0.1.52 as implemented in the FragPipe DIA workflow (Demichev et al., 2022; Yu et al., 2023)

All 12 diaPASEF runs were searched together so that the empirical spectral library, peptide-level FDR estimation, and retention-time and ion-mobility calibration were derived from the full acquired dataset. Four runs belonged to an additional genotype that is not included in the comparison reported here; these samples contributed to library construction and identification only and were excluded before differential testing.

##### DIA quantification with DIA-NN and match-between-runs

Because the FragPipe DIA_SpecLib_Quant_diaPASEF workflow calls DIA-NN without the -- reanalyse flag, cross-run identity propagation in that workflow occurs only at the level of the shared empirical library. To additionally use DIA-NN two-pass match-between-runs (MBR) re-extraction, all 12 diaPASEF runs were reprocessed with DIA-NN v2.2.0 Academia (Demichev et al., 2020; Demichev et al., 2022) from the command line against the FragPipe-derived empirical library, with MBR enabled (--reanalyse) and precursor- and protein-matrix q-value thresholds of 0.01 (--qvalue 0.01 --matrix-qvalue 0.01). In this two-pass procedure, a refined library is assembled from confidently identified precursors pooled across runs and each run is then re-extracted against that library; no value is written where no matching chromatographic signal is present. An otherwise identical DIA-NN run with MBR disabled was retained as a sensitivity check. The precursor-level report (report.pr_matrix.tsv) and MaxLFQ protein-group report (report.pg_matrix.tsv; (Cox et al., 2014)) were used for downstream analyses.

##### Experimental design and sample assignment

The comparison reported here comprised eight samples: four biological replicates of WT and four of *pamiRlhcb1-s*. A further four samples of an additional genotype were acquired in the same experimental series and included in the database search and DIA-NN quantification described above, but were removed from the data matrices before filtering, normalization, imputation, or statistical testing; no result is reported for these samples.

##### Proteomics statistical analysis

All proteomics statistical analyses were performed in R v4.5.0 with DEP2 v0.3.7.3 through its Shiny interface (Feng et al., 2023). Two complementary analyses were run on the same DIA-NN output. In both analyses, the four additional samples were removed from the imported matrix before any processing step, so that filtering thresholds, normalization parameters, and imputation models were estimated only from the eight samples entering the *pamiRlhcb1-s* versus WT comparison.

##### (i) Genome-wide differential abundance at protein level

The DIA-NN protein-group matrix was imported into DEP2 using Protein.Group as the unique identifier and was log_2_-transformed on import. LHCB isoforms analyzed separately below were excluded from this matrix so that the genome-wide analysis describes the remaining proteome and is not driven by the directly targeted antenna proteins. Protein groups were retained when quantified in at least three replicates of at least one condition, normalized by variance-stabilizing normalization (vsn; (Huber et al., 2002)), and remaining missing values were imputed by Bayesian principal-component analysis (bPCA; pcaMethods; (Stacklies et al., 2007)), which models values missing at random rather than left-censored dropout. Differential abundance was tested with the limma-based empirical-Bayes linear model (Liu et al., 2015) for *pamiRlhcb1-s* versus WT. Proteins were considered differentially abundant at Benjamini-Hochberg-adjusted P < 0.05 and |log_2_ fold change| > 1.

##### (ii) Isoform-resolved quantification of LHCB proteins

Because LHCB1 and LHCB2 isoforms are 95-100% identical within their subfamilies, protein-group-level MaxLFQ values contain shared-peptide information and cannot by themselves resolve individual isoforms. Isoform-level abundance changes were therefore derived from unique DIA precursors.

The complete DIA-NN precursor matrix was imported into DEP2 using Precursor.Id (encoding sequence, modifications, and charge) as the unique identifier and was log2-transformed on import. The complete precursor matrix, rather than a pre-selected LHCB subset, was carried through filtering, normalization, and imputation. Precursors were retained when quantified in at least one replicate of at least one condition so that precursors showing complete dropout in one genotype were not removed. Missing values were imputed with QRILC (quantile regression imputation of left-censored data; (Lazar et al., 2016)), consistent with missing-not-at-random (MNAR) dropout below the detection limit. The use of bPCA for the near-complete protein-group matrix and QRILC for the more strongly left-censored precursor matrix was deliberate and reflects their different missingness patterns. DEP2 per-precursor differential tests were not used for the isoform-level inference.

Unique precursors were selected using the DIA-NN “Proteotypic” flag. For each precursor, the mean log_2_ intensity across the four WT replicates was subtracted from the precursor value in every sample, yielding a log_2_ ratio relative to WT and removing the arbitrary between-precursor intensity offset. These centered values from all precursors assigned to an isoform were averaged within each biological replicate. Genotype effects were tested with t-tests with Welch correction on these replicate-level isoform values against WT, and the difference between group means is reported as the log_2_ fold change.

### Immunoblot analysis

Total leaf protein was extracted as described previously (Brandt et al., 2026). Leaf material was disrupted directly in precooled extraction buffer (50 mM Tris, pH 7.5, 150 mM NaCl, 2.5 mM EDTA, 10% glycerol, 0.5% NP-40, 5 mM DTT, and 1× plant protease inhibitor cocktail) using a plastic pestle; alternatively, frozen ground tissue was resuspended in the same buffer. Cell debris was removed by centrifugation at 25,000 g for 10 min at 4°C. Protein extracts were separated by SDS-PAGE and transferred to PVDF membranes by wet blotting. Membranes were blocked for 1 h at room temperature in 5% milk powder in TBS-T (0.05% Tween-20), incubated with primary antibody overnight at 4°C, washed in TBS-T, incubated with secondary antibody for 45 min at room temperature, washed again, and visualized with homemade ECL using an ImageQuant system (GE Healthcare). TIFF images were imported into Fiji and adjusted only at the level of the complete blot before export. Loading was normalized by colorimetric analysis of total band intensities above RbcL and controlled by Coomassie staining of the same membrane. KEA1 and KEA2 was detected with the N-terminal KEA1/2 antibody (Bölter et al., 2020). The antibody detects both KEA1 and KEA2.

### Blue-native PAGE of thylakoid complexes

BN-PAGE was performed as described previously (Nickel et al., 2016): enriched chloroplasts were used immediately after isolation and chlorophyll concentration was determined spectrophotometrically after extraction in 80% acetone. For the analytical BN-PAGE, loading was adjusted indirectly to approximate equal fresh-weight equivalents. *pamiRlhcb1-s* contained approximately 70% of the WT chlorophyll content on a fresh-weight basis. Therefore, 15 µg chlorophyll was loaded for WT and 10.5 µg chlorophyll for *pamiRlhcb1-s*. Chloroplasts were pelleted at 500 g for 3 min at 4°C and resuspended in ACA buffer (50 mM Bis-Tris, pH 7.5, 750 mM aminocaproic acid, 0.5 mM EDTA). Thylakoid complexes were solubilized with dodecyl maltoside at a final concentration of 0.5% (w/v) for 25 min on ice and clarified at 16,000 g at 4°C. BN loading dye was added to final concentrations of 68 mM aminocaproic acid and 0.45% (w/v) Coomassie Brilliant Blue G-250, and samples were separated on 4-12% acrylamide gels. Gels were run with blue cathode buffer (50 mM Tricine, 15 mM Bis-Tris, 0.2% Coomassie Brilliant Blue G-250) and 50 mM Bis-Tris (pH 7.0) anode buffer; after the dye front reached approximately two-thirds of the gel, the cathode buffer was replaced by clear cathode buffer. Four biological replicates per genotype are shown.

### PAM analysis

PAM measurements outside screen I were measured on 2-3 weeks old and dark adapted plants (15 min) plants using the IMAGING-PAM (WALZ). For the *pamiRlhcb1-s* segregation experiment, F_0_ was used as the predefined quantitative phenotype. WT mean and standard deviation were calculated from the control population, and F2 plants with F_0_ below the WT mean minus two standard deviations were classified as *pamiRlhcb1-s*-like.

### ABA root-elongation assay

Seeds were surface sterilized, stratified for 3 d, germinated on half-strength MS medium, and grown vertically for 7 d. Seedlings were then transferred to plates containing solvent control (Mock), 5 µM +/− ABA, or 10 µM +/− ABA. Root tips were marked at transfer and root elongation was measured over the following 4 d. WT, *aba1-6*, and *pamiRnced-s* were each distributed across three plates to minimize plate-dependent effects. All plates for an experiment were prepared in one batch from the same base MS medium, ABA or solvent was added subsequently and identical medium volumes were poured per plate. For relative-root-growth plots, ABA-treated root lengths were normalized to the corresponding mean mock-grown value. Genotype and time effects were tested by two-way ANOVA followed by Sidak multiple-comparisons correction.

### Use of artificial intelligence

Generative artificial intelligence tools were used during manuscript preparation and revision. ChatGPT (OpenAI) was used to assist with language editing and text structuring. The automized WMD3 re-query was performed with assistance from Claude Science (Anthropic). All AI-assisted outputs, including re-query results and derived output tables, were critically checked and verified by the authors against the underlying data and relevant literature. AI was not used to generate experimental data, and all scientific interpretation and conclusions remain the responsibility of the authors.

## Supporting information

Supplementary Dataset S1 - pamiR_master

Supplementary Dataset S2 - WMD3 alignments

Supplementary Dataset S3 - Results pamiRlhcb1-s proteomics

Supplementary File S1 - pamiR handling instructions

Supplementary File S2 - Example pamiR vector

## Acknowledgements

We are grateful for technical undergrad assistance by LMU alumni Annika Schnappinger. Thanks to Drs. Bettina Bölter and Andreas Brachmann (both LMU Munich) for critical reading of the manuscript and for help with NGS, respectively. Chuck Cody and Albert Schorer, gardeners at WSU and LMU, respectively, provided tremendous support for the project. Dr. Rachael A. DeTar (WSU, LMU, now at CSU) helped with NGS analysis of a pilot library. Thanks to Dr. Kenneth Chang (CSHL) for initial project consultation.

## Author Contributions

H.-H.K. designed the project. H.-H.K and B.B. wrote the paper. B.B. built the *pamiR* library, carried out NGS quality control, designed and prepared figures, and wrote the manual. A.I.P. built an earlier version of the library. A.I.P., D.S., C.M.L, C.L.L, C.E., D.S, S.S. and B.B. carried out proof of concept experiments. F.H. designed multi-gene *pamiR* oligos. R.S. provided *in silico* plastid target candidate lists from ARAMEMNON. All coauthors contributed to the editing. H.-H.K. is responsible for contact and ensuring communication.

## Data availability

The author responsible for distribution of materials integral to the findings presented in this article in accordance with the policy described in the Instructions for Authors is Hans-Henning Kunz. Next-generation sequencing data are deposited with the European Nucleotide Archive. The *pamiR* plasmid library can be ordered from Addgene and the European Plasmid Archive. Mass-spectrometry raw data and search results are deposited to the ProteomeXchange Consortium via the PRIDE partner repository (Perez-Riverol et al., 2021).

## Supplementary Figure Legends

**Supplementary Figure S1. Plastid enrichment and *pamiR* design networks.** A) Relative localization composition of the curated *pamiR* target set and the WT proteome based on the SUBA/MMAP analysis, grouped as plastid, unassigned, or other organelles. B) Network atlas of designed *pamiR*-gene relationships. Target genes are shown as yellow rectangles, *pamiR*s as blue ellipses, and gray edges connect each *pamiR* to its designed target gene(s). The networks illustrate whole family or co-targetable-subset coverage: only networks with more than three nodes are shown. Networks depict design relationships rather than measured silencing.

**Supplementary Figure S2. *pamiR* cloning workflow and pooled-library quality control.** A) Multistep cloning workflow showing addition of cloning overhangs, insertion of *pamiR* oligonucleotides and the MIR319a stem loop by sequential Golden Gate reactions, and removal of residual starting vector by restriction/ligation. B) Bioanalyzer trace following diagnostic digestion of the final *pamiR* pool. Peaks corresponding to constructs without (331 bp) and with (413 bp) the inserted stem loop are indicated. Areas under the respective peaks were used to estimate >97% stem-loop integration. C) Representative stringent mapping of NGS reads across the variable amiR*/amiR regions and invariant backbone sequence. Blue lines indicate exact matches, white boxes missing bases, gray boxes an ambiguous base call, and green boxes a T mismatch.

**Supplementary Figure S3. Recommended workflow for identification and independent validation of *pamiR* screen hits.** Screening yields candidate plants, after which the *pamiR* guide is identified by PCR and Sanger sequencing. Complementary validation routes include testing persistence of the phenotype in the next generation, using an independent amiRNA targeting the same gene(s), comparing an independent loss-of-function line generated by T-DNA insertion or CRISPR/Cas, backcrossing to WT, and directly verifying depletion of the targeted protein(s) by immunoblotting or proteomics. These approaches are complementary rather than a mandatory linear sequence.

**Supplementary Figure S4. Additional *pamiR* lines and quantitative phenotyping from proof-of-concept screen 1.** A) T_2_ *pamiR* lines with visible and chlorophyll-fluorescence images; columns show visible RGB images, *F*_v_/*F*_m_, steady-state NPQ, and steady-state Y(II) recorded at 185 PAR. “-s” denotes a guide designed to target more than two loci. B-G) Mean growth rate, hue variability, *F*_v_/*F*_m_, Y(II), Y(NPQ), and Y(NO), respectively. H) NPQ induction kinetics recorded at 185 PAR. Data are presented as mean ± SD with n > 8 biological replicates per genotype. For B-G, each *pamiR* line was compared independently with WT using an unpaired two-tailed t-test with Welch’s correction. Asterisks under the bar graphs indicate significant differences from WT: *P < 0.05, **P < 0.01, ***P < 0.001.

**Supplementary Figure S5. Additional BN-PAGE replicates and absolute root-elongation data.** A) BN-PAGE of thylakoid complexes from WT and *pamiRlhcb1-s* biological replicates 3 and 4, complementing replicates 1 and 2 shown in Fig. 4D. Loading was adjusted as in Fig. 4D to approximate equal fresh-weight equivalents. Regions corresponding to LHCII trimers, higher-order LHCII assemblies, cytochrome *b*_6_*f*, and PSII monomers are indicated. B-D) Absolute root lengths of WT, *pamiRnced-s*, and *aba1-6* for 4 d after transfer to mock medium (B), 5 µM ABA (C), or 10 µM ABA (D); Mean +/−SD in n ≥ 8 biological replicates per genotype. These are the absolute measurements underlying the normalized analyses in Fig. 5C and D.

## Supplementary Data

### Supplementary Figures and Tables

**Supplementary Figure S1.** Plastid enrichment and *pamiR* design networks.

**Supplementary Figure S2.** *pamiR* cloning workflow and pooled-library quality control.

**Supplementary Figure S3.** Recommended workflow for identification and independent validation of *pamiR* screen hits.

**Supplementary Figure S4.** Additional *pamiR* lines and quantitative phenotyping from proof-of-concept screen 1.

**Supplementary Figure S5.** Additional BN-PAGE replicates and absolute root-elongation data.

**Supplementary Table S1.** Overview of isolated *pamiR* mutants.

**Supplementary Table S2.** Resolution of LHCB isoforms in the *pamiRlhcb1-s* DIA proteomics: peptide and unique-peptide counts, whether each isoform is individually quantifiable, and whether it is targeted by *pamiR-983*.

**Supplementary Table S3.** Alignment characteristics of *pamiR-1034* across the nine CCD/NCED family members.

**Supplementary Table S4.** Criteria, rationale, and distribution of *pamiR* off-target specificity classes.

**Supplementary Table S5.** DNA primers used for library cloning, NGS, and guide identification.

### Supplementary Datasets

**Supplementary Dataset S1.** *pamiR* master table. Guide sequences, intended-target annotation, WMD3 re-query summary, per-construct off-target prioritization, and NGS read counts for all 2,285 *pamiR* constructs.

**Supplementary Dataset S2.** WMD3 alignment dataset. All 4,862 alignments recovered in the stated WMD3 re-query, including mismatch positions and relative hybridization-energy information.

**Supplementary Dataset S3.** *pamiRlhcb1-s* proteomics dataset. Protein-group differential-abundance results for the *pamiRlhcb1-s* versus WT LFQ analysis.

### Supplementary Files

**Supplementary File S1.** *pamiR* handling instructions.

**Supplementary File S2.** Example *pamiR* vector (pG2-FR_*pamiR*).

