## Supplemental Figures for "pamiR: INVESTIGATING PLANT CELLS ONE ORGANELLE AT A TIME"

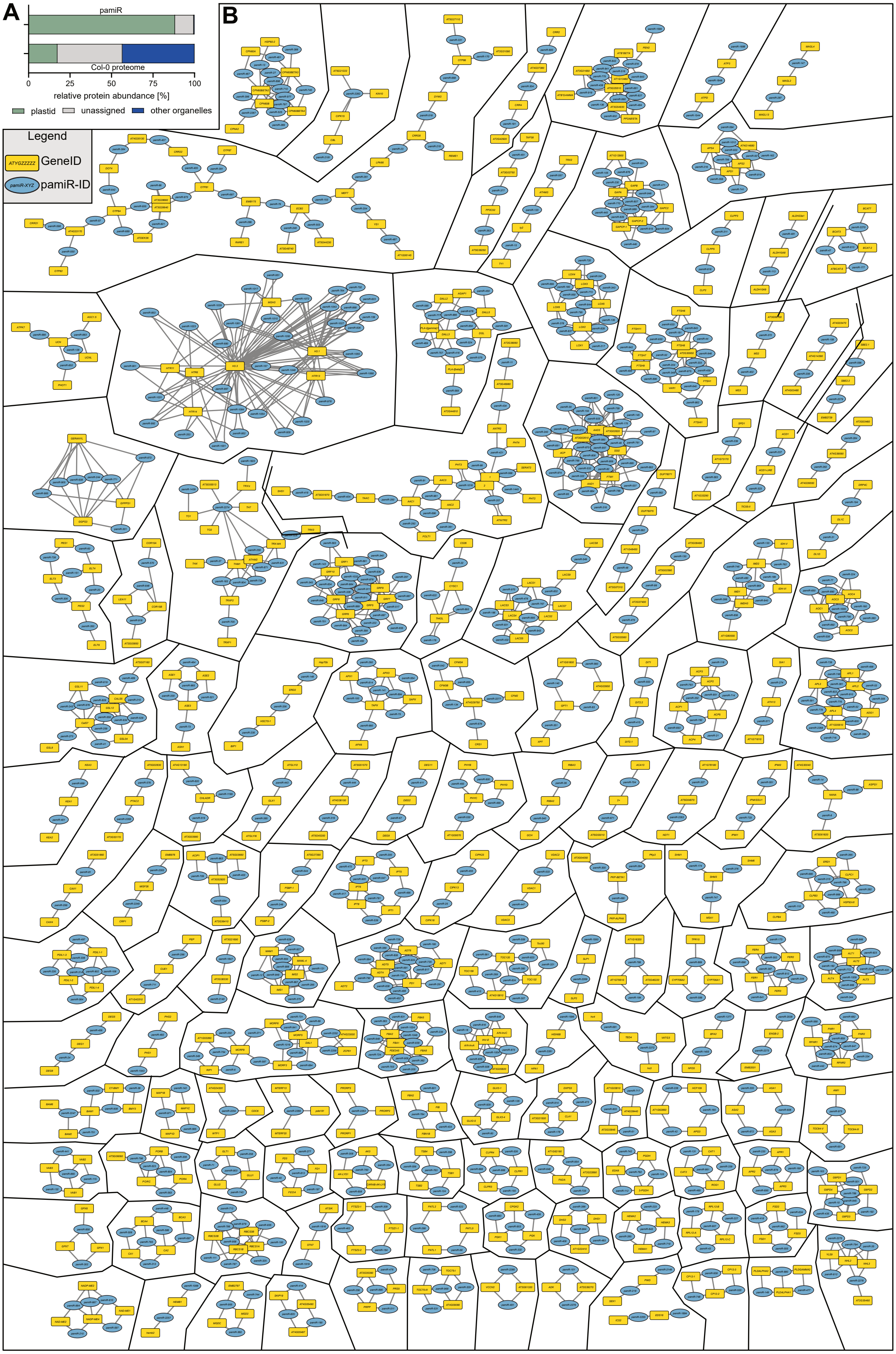


**Supplementary Figure S1. Plastid enrichment and *pamiR* design networks.**

A) Relative localization composition of the curated *pamiR* target set and the WT proteome based on the SUBA/MMAP analysis, grouped as plastid, unassigned, or other organelles. B) Network atlas of designed *pamiR*-gene relationships. Target genes are shown as yellow rectangles, *pamiR*s as blue ellipses, and gray edges connect each *pamiR* to its designed target gene(s). The networks illustrate whole family or co-targetable-subset coverage: only networks with more than three nodes are shown. Networks depict design relationships rather than measured silencing.

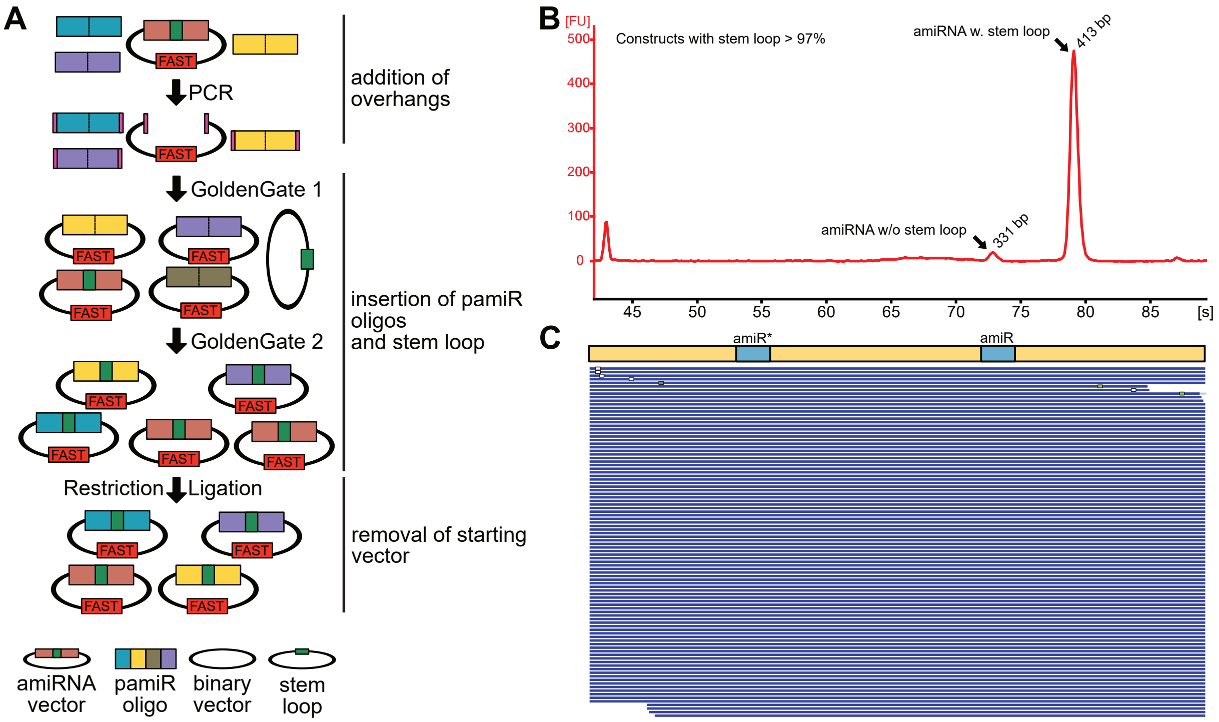


**Supplementary Figure S2. *pamiR* cloning workflow and pooled-library quality control.**

A) Multistep cloning workflow showing addition of cloning overhangs, insertion of *pamiR* oligonucleotides and the MIR319a stem loop by sequential Golden Gate reactions, and removal of residual starting vector by restriction/ligation. B) Bioanalyzer trace following diagnostic digestion of the final *pamiR* pool. Peaks corresponding to constructs without (331 bp) and with (413 bp) the inserted stem loop are indicated. Areas under the respective peaks were used to estimate >97% stem-loop integration. C) Representative stringent mapping of NGS reads across the variable amiR*/amiR regions and invariant backbone sequence. Blue lines indicate exact matches, white boxes missing bases, gray boxes an ambiguous base call, and green boxes a T mismatch.

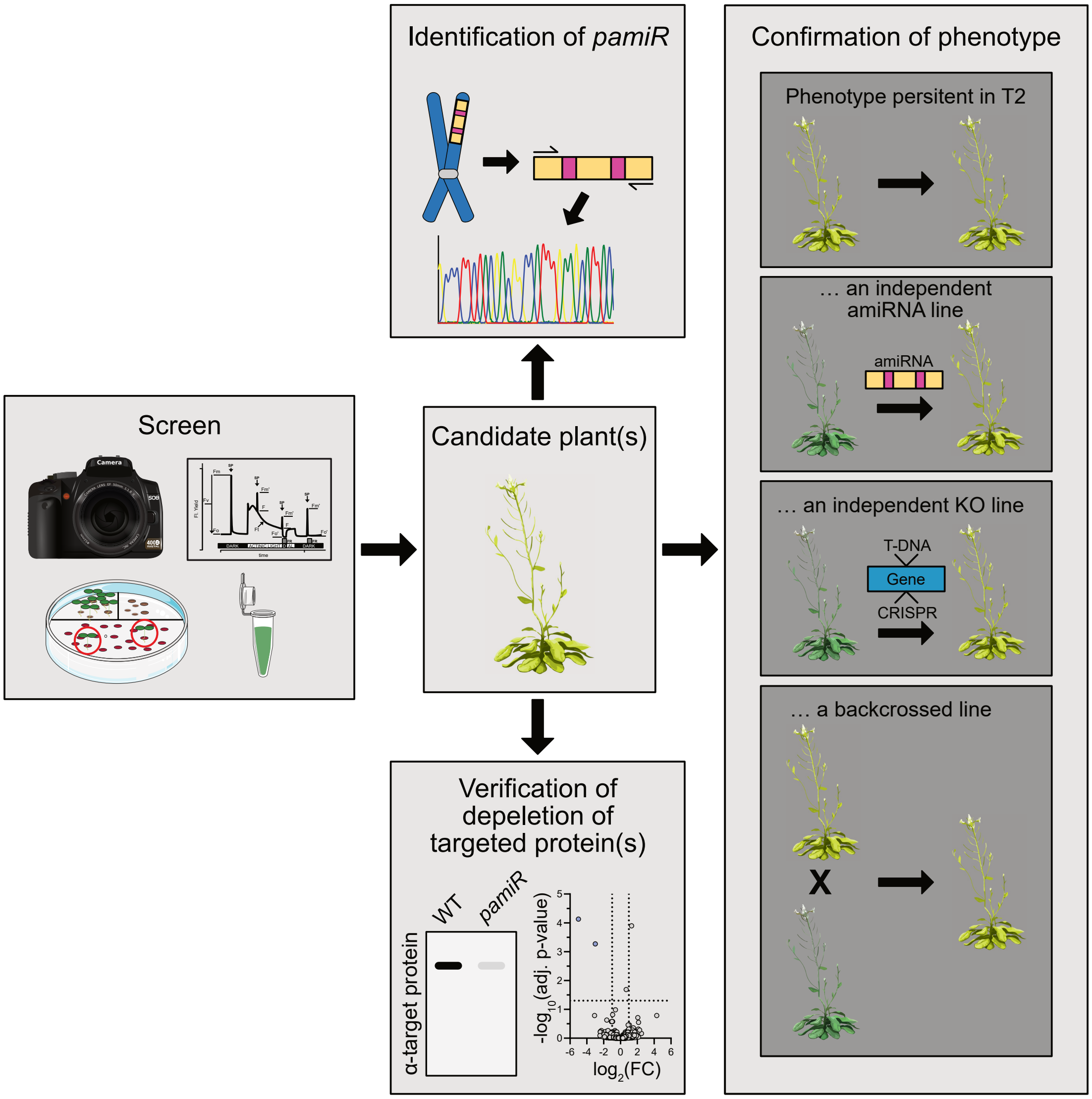


**Supplementary Figure S3. Recommended workflow for identification and independent validation of *pamiR* screen hits.**

Screening yields candidate plants, after which the *pamiR* guide is identified by PCR and Sanger sequencing. Complementary validation routes include testing persistence of the phenotype in the next generation, using an independent amiRNA targeting the same gene(s), comparing an independent loss-of-function line generated by T-DNA insertion or CRISPR/Cas, backcrossing to WT, and directly verifying depletion of the targeted protein(s) by immunoblotting or proteomics. These approaches are complementary rather than a mandatory linear sequence.

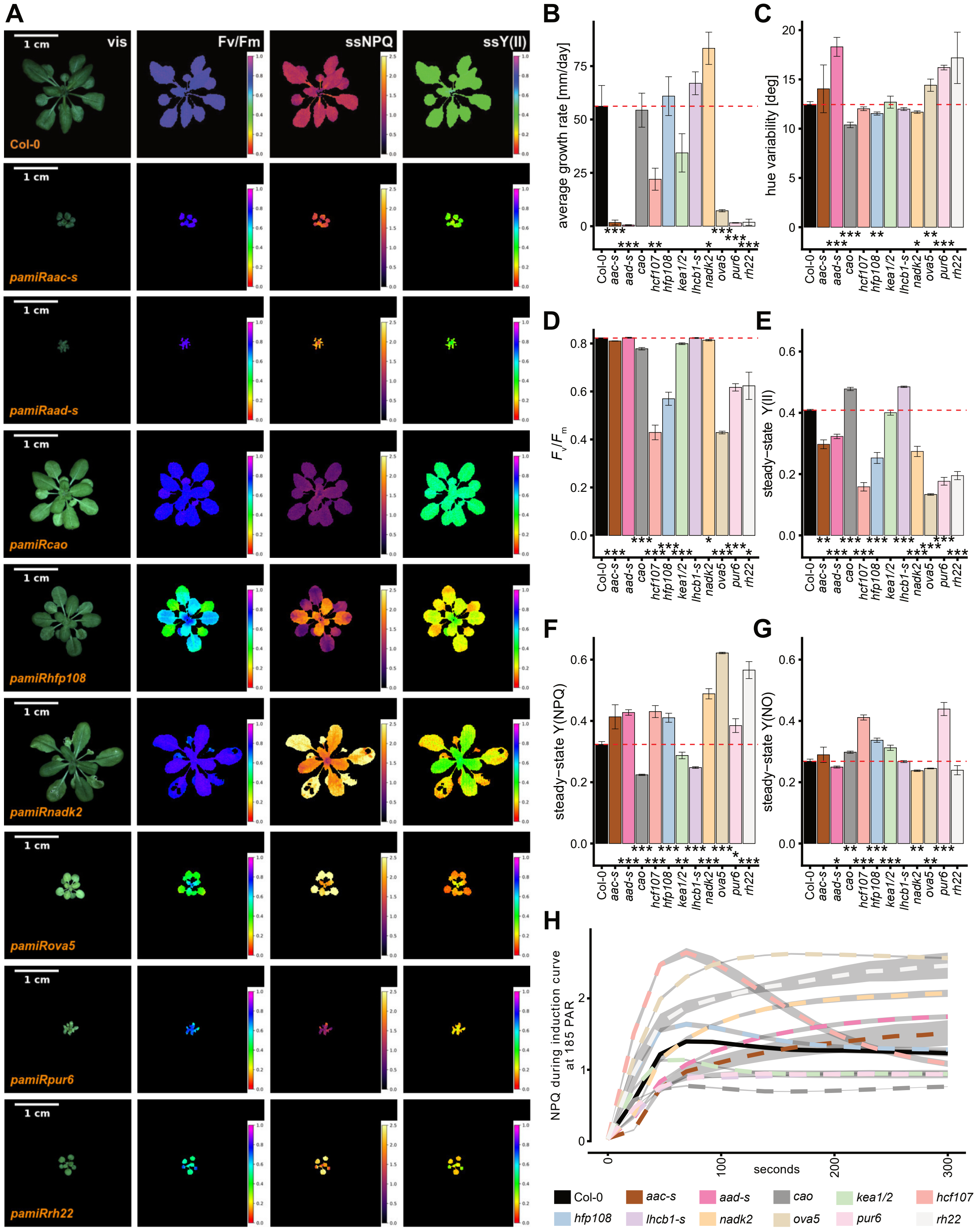


**Supplementary Figure S4. Additional *pamiR* lines and quantitative phenotyping from proof-of-concept screen 1.** A) $T_2$ *pamiR* lines with visible and chlorophyll-fluorescence images; columns show visible RGB images, $F_v/F_m$, steady-state NPQ, and steady-state Y(II) recorded at 185 PAR. “-s” denotes a guide designed to target more than two loci. B-G) Mean growth rate, hue variability, $F_v/F_m$, Y(II), Y(NPQ), and Y(NO), respectively. H) NPQ induction kinetics recorded at 185 PAR. Data are presented as mean ± SD with n > 8 biological replicates per genotype. For B-G, each *pamiR* line was compared independently with WT using an unpaired two-tailed t-test with Welch’s correction. Asterisks under the bar graphs indicate significant differences from WT: *P < 0.05, **P < 0.01, ***P < 0.001.

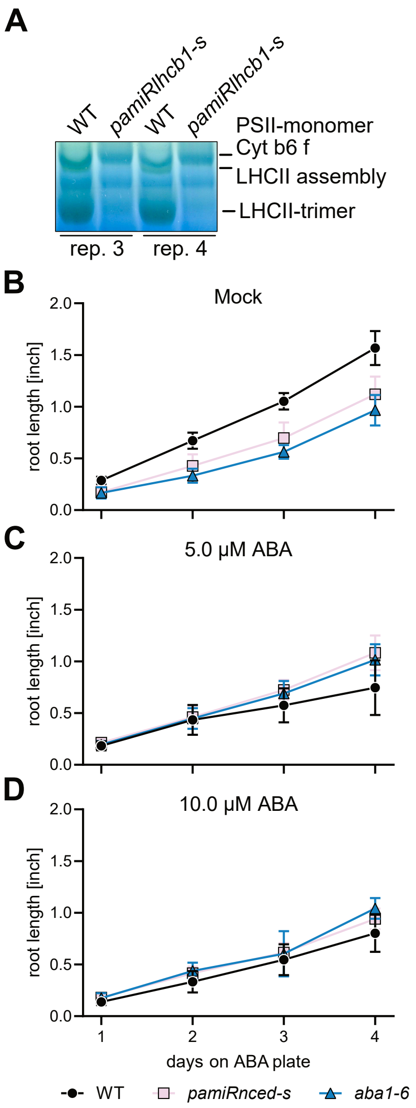


**Supplementary Figure S5. Additional BN-PAGE replicates and absolute root-elongation data.**

A) BN-PAGE of thylakoid complexes from WT and *pamiRlhcb1-s* biological replicates 3 and 4, complementing replicates 1 and 2 shown in Fig. 4D. Loading was adjusted as in Fig. 4D to approximate equal fresh-weight equivalents. Regions corresponding to LHCII trimers, higher-order LHCII assemblies, cytochrome $b_6f$, and PSII monomers are indicated. B-D) Absolute root lengths of WT, *pamiRnced-s*, and *aba1-6* for 4 d after transfer to mock medium (B), 5 µM ABA (C), or 10 µM ABA (D); Mean +/-SD in n ≥ 8 biological replicates per genotype. These are the absolute measurements underlying the normalized analyses in Fig. 5C and
