## Supplemental Tables for "pamiR: INVESTIGATING PLANT CELLS ONE ORGANELLE AT A TIME"

Supplementary Table S1 - Overview of isolated mutants

| Mutant | AGI(s) | Functional Annotation | Growth | $F_v/F_m$ | $\phi$II | NPQ | Chlor-osis | amiR sequence/pamiR-ID | Ref. (DOI) |
|---|---|---|---|---|---|---|---|---|---|
| ***pamiRhcf107*** | AT3G17040 | tetratricopeptide repeat-containing protein | ↓ | ↓ | ↓ | ↑ | **x** | TTACATCGAGGTTAAACGCGC / pamiR-1990 | 10.1111/tpj.12632 |
| ***pamiRkea1/2*** | AT1G01790<br>AT4G00630 | $K^+$ efflux antiporter | | ↓ | | ↓ | **x** | TTATACCCTGATTTACGGCTT/ pamiR-401 | 10.1073/pnas.1323899<br>111 |
| ***pamiRrh22*** | AT1G59990 | RNA helicase | ↓ | ↓ | ↓ | ↑ | **x** | TAAATATACTGCTTACTGCGG / pamiR-1157 | 10.1104/pp.111.18677<br>5 |
| ***pamiRnadk2*** | AT1G21640 | $NAD^+$ kinase | ↑ | ↓ | ↓ | ↑ | | TAAAATTCTACAATGCGCCTC / pamiR-1108 | 10.1007/s11103-005-<br>6802-y |
| ***pamiRaac-s*** | AT3G08580<br>AT5G01500 | ADP/ATP carrier | ↓ | ↓ | ↓ | | | TTCCACGGTACAGGCCAGCAC / pamiR-250 | 10.1074/jbc.M6091302<br>00 |
| ***pamiRhfp108*** | AT3G63190 | ribosome recycling factor | | ↓ | ↓ | ↑ | | TACTAGACTAGAAAAGACCTG / pamiR-1441 | 10.1007/s11103-010-<br>9653-0 |
| ***pamiRcao*** | AT1G44446 | chlorophyll a oxygenase | | ↓ | ↑ | ↓ | **x** | TAAGTGTGGACTATAAGCCGG / pamiR-1298 | 10.1093/mp/ssp117 |
| ***pamiRpur6*** | AT2G37690 | aminoimidazole ribonucleotide carboxylase | ↓ | ↓ | ↓ | ↑ | **x** | TAAACTTAAAGCGGATGGCGA / pamiR-1132 | n/a |
| ***pamiRova5*** | AT3G13490 | lysyl-tRNA aminoacylation | ↓ | ↓ | ↓ | ↑ | **x** | TTATCAACATACCGTTGGCCG / pamiR-2060 | 10.1111/j.1365-<br>313X.2007.03076.x |
| ***pamiRaad-s*** | AT2G43710<br>AT3G02610<br>AT3G02630 | stearoyl-ACP desaturase | ↓ | | ↓ | ↑ | | TATGACGCTTCTCGTCCGCAC / pamiR-972 | 10.1007/s0012200505<br>04 |
| ***pamiRlhcb1-s*** | AT1G29910<br>AT1G29920<br>AT1G29930<br>AT2G34420 | light harvesting complex of PSII | | | ↑ | ↓ | | TTTGTCACACGGCCGCTCCCT / pamiR-983 | 10.1105/tpc.114.12737<br>3 |
| ***pamiRnced-s*** | AT1G30100<br>AT1G78390<br>AT3G14440<br>AT3G24220<br>AT4G18350 | nine-cis-epoxycarotenoid dioxygenase | | | | | | TCAAACCGGCGTTAGTTACAC / pamiR-1034 | 10.1111/j.1365-<br>313X.2005.02622.x |

Supplementary Table S2 - Overview of resolution of *pamiRlhcb1-s* proteomics

| Gene/isoform | TAIR/Araport locus | # peptides | # unique peptides | Quantifiable | Targeted by *pamiRlhcb1-s* |
|---|---|---|---|---|---|
| **LHCB1.1** | AT1G29920 | 0 | 0 | FALSE | TRUE |
| **LHCB1.2** | AT1G29910 | 6 | 0 | FALSE | TRUE |
| **LHCB1.3** | AT1G29930 | 1 | 1 | TRUE | TRUE |
| **LHCB1.4** | AT2G34430 | 11 | 4 | TRUE | FALSE |
| **LHCB1.5** | AT2G34420 | 3 | 2 | TRUE | TRUE |
| **LHCB2.1** | AT2G05100 | 4 | 0 | FALSE | FALSE |
| **LHCB2.2** | AT2G05070 | 0 | 0 | FALSE | FALSE |
| **LHCB2.3 (=LHCB2.4)** | AT3G27690 | 0 | 0 | FALSE | FALSE |
| **LHCB3** | AT5G54270 | 6 | 5 | TRUE | FALSE |

Supplementary Table S3 - Alignment characteristics of *pamiRnced-s*

| Gene | Tair GeneID | # Mismatch | Guide_MM_Positions | WMD3_Pct | Retrieved_By_WMD3 | Mm_In_2_12 | Mm_At_Cleavage |
|---|---|---|---|---|---|---|---|
| **NCED3** | AT3G14440 | 2 | 1;16 | 98.26 | TRUE | 0 | FALSE |
| **NCED6** | AT3G24220 | 4 | 1;4;16;20 | 86.17 | TRUE | 1 | FALSE |
| **NCED2** | AT4G18350 | 4 | 1;14;16;20 | 84.52 | TRUE | 0 | FALSE |
| **NCED5** | AT1G30100 | 4 | 1;16;17;20 | 82.39 | TRUE | 0 | FALSE |
| **NCED9** | AT1G78390 | 4 | 1;2;16;17 | 73.04 | TRUE | 1 | FALSE |
| **CCD4/NCED4** | AT4G19170 | 7 | 6;9;12;14;15;18;21 | | FALSE | 3 | FALSE |
| **MAX3/CCD7** | AT2G44990 | 8 | 1;2;10;13;14;16;19;21 | | FALSE | 2 | TRUE |
| **MAX4/CCD8** | AT4G32810 | 8 | 2;4;5;6;8;11;12;14 | | FALSE | 7 | TRUE |
| **CCD1** | AT3G63520 | 9 | 8;10;12;15;16;17;19;20;21 | | FALSE | 3 | TRUE |

Supplementary Table S4 – *pamiR* off-target specificity classes

| Risk Class | Label | Criterion | Rationale | # Off-Target Alignments | % Off-Target Alignments | # of *pamiRs* | % of *pamiRs* |
|---|---|---|---|---|---|---|---|
| **A** | high | no mismatch at position 10 or 11 **and** ≥1 mismatch in positions 2-12 **and** ≥75 % of perfect-match hybridization energy | duplex is cleavage-competent **and** thermodynamically strong enough to be loaded/retained - the only class where silencing is plausible | 98 | 9.5 | 95 | 4.2 |
| **B** | moderate | mismatch pattern as in class A, but energy <75 % of perfect match | geometry permits slicing, but binding is weaker than any validated amiRNA-target duplex in the library | 496 | 48.2 | 380 | 16.6 |
| **C** | low | ≥2 mismatches within positions 2-12 (the WMD specificity region) | the criterion WMD itself uses to reject a target; two central/5' mismatches strongly impair slicing | 224 | 21.8 | 139 | 6.1 |
| **D** | negligible | mismatch at position 10 or 11 | plant AGO1 cuts between the bases pairing guide positions 10 and 11; a mismatch there abolishes cleavage | 210 | 20.4 | 107 | 4.7 |
| **E** | none | no predicted off-target(s) at WMD3 defaults | nothing outside the declared target set is returned | | | 1564 | 68.4 |

Supplementary Table S5 – DNA primers used

| Prime Name | Primer Sequence 5' -> 3' |
|---|---|
| **GG_lib_f** | GGCTACGGTCTCTTAATACTCGCTGTTTTGAATTGATG |
| **GG_lib_r** | GGCTACGGTCTCAAAAGATTAATCAAGAAAATTGGAATAC |
| **GG_v_f** | GGCTACGGTCTCTATTAAGTGTATGAACATGTGTAATATG |
| **GG_v_r** | GGCTACGGTCTCACTTTCCTGCACAAAAAC |
| **P5_NGS_f** | TACACGACGCTCTTCCGATCTGCTCGGACGCATATTAC |
| **P7_NGS_r** | CAGACGTGTGCTCTTCCGATCTGGCAGCATATATGTCACTTAG |
| **mir319_f** | CAAACACACGCTCGGAC |
| **HSP18_r** | CACAAATTCATAACACAACAAGC |
