## Supplementary File S1 - pamiR handling instructions for "pamiR: INVESTIGATING PLANT CELLS ONE ORGANELLE AT A TIME"

### 1 Handling of the plastid amiRNA library (*pamiR*)

Introduction:

The *pamiR* consists of ~2300 plasmids in a modified pGreenII vector backbone (Hellens et al., 2000; Schwartz et al., 2025) carrying a Kanamycin resistance cassette for selection in bacteria. Successful amplification of the pGreen vector in Agrobacterium requires the presence of the helper plasmid pSOUP (Hellens et al., 2000).

In order to maintain the complexity of the library and to ensure that all plant mutant screens use the maximum of available constructs, it is mandatory to always have a sufficient number of colonies/cells. Please adhere to the instructions below to make sure of that.

#### 1.1 Amplification of *pamiR* from plasmid stocks and preparation of high-density glycerol stocks

Notes:

To maintain the complexity and distribution of constructs, re-transformation should be performed at a minimum of 30× coverage of the plasmid pool. This corresponds to more than 70000 colony-forming units (CFUs) during library amplification. Coverage can be verified either by plating a small aliquot of the transformation mixture or by plating the entire transformation on LB-agar plates under antibiotics selection.

Although the plasmid size within the library is uniform, amplification biases can still arise during liquid culture. Therefore, the protocol below minimizes growth in liquid culture and includes a CFU assessment step prior to the preparation of amplified plasmid DNA and glycerol stocks.

It is recommended to transform the starting plasmid material only once, generate sufficient plasmid DNA for most downstream applications, and prepare glycerol stocks directly from this initial transformation. Repeated rounds of transformation and amplification are discouraged.

Transformation can be performed using either chemically competent or electrocompetent cells. Electroporation generally yields higher transformation efficiencies but is more sensitive to residual salts in the DNA sample. Chemical transformation of E. coli is more robust, although typically less efficient. Below, we provide a standard protocol for chemical transformation. Users should adapt the highest-efficiency protocol recommended by the manufacturer of the competent cells.

As a starting point, we recommend performing four parallel transformations and plating them onto four large LB-agar plates (15 cm diameter).

Supplies needed:

- *pamiR* plasmid stock: use at least 100 ng per transformation
- Highly chemically or electro-competent E. coli cloning strains, we use Takara Stellar
- Sterile 2 ml Eppendorf tubes
- SOC or LB medium
- LB-Agar big petri dishes (15 cm diameter) supplemented with Kanamycin (50 µg/ml) → LB-Agar K

- 500 ml of liquid LB supplemented with Kanamycin (50 µg/ml) in Erlenmeyer flask → LB-K
- Sterile serological pipette
- Sterile centrifugation containers
- Sterile 60% Glycerol in LB-medium
- Liquid nitrogen

Instructions:

1. Pre-heat SOC or LB medium and LB-Agar K at 37°C
2. All steps should be carried out near a flame or under a sterile hood
3. Thaw competent cells on ice (50-100 µl in 1.5 ml Eppendorf tube)
4. Add at least 100 ng but not more than 1 µl plasmid/10 µl of bacteria per transformation
5. Store on ice for 30 min
6. Heat shock at 42°C for 60 sec
7. Immediately place back on ice and store for 3 min
8. Add 1 ml or pre-heated SOC or LB to competent cells to each tube
9. Incubate at 37°C for 60 min
10. Plate content of tube(s) on LB-Agar K and incubate plates over night (o/N) at 37°C
11. Check plate(s) the next day. When in doubt, count cells on a fraction of the plate. Plate(s) should have more than 70000 colonies
12. Add ~20 ml LB (KAN) directly onto the plate(s) and re-suspend bacteria using a cell spreader or by careful up-and-down pipetting
13. Combine all responded cell suspensions in Erlenmeyer containing the LB-K and incubate at 37°C for 90 min
14. Remove culture for the plasmid preparation and pellet cells (4000*g* for 5 min). These pellets can be stored at -20°C for long term or at 4°C for a few hours. Carry out plasmid preparation according to the manufacturer's instruction. Safe 220 ml for the preparation of high-density glycerol stocks as described below
15. Pellet cells from 200 ml by centrifugation (e.g., 2 x 100 ml in 2 x 50 ml falcon tubes) (4000*g* for 5 min)
16. Resuspend pellet(s) in 7.5 ml cell culture and add 7.5 ml sterile 60% Glycerol in LB-medium
17. Aliquot into 2 ml Eppendorf tubes and flash freeze in liquid nitrogen
18. Store at -80°C
19. Use an entire aliquot of glycerol stock for the amplification

#### 1.2 Transformation of *pamiR* into Agrobacteria and preparation of high-density glycerol stocks

Notes:

To preserve the complexity and distribution of constructs, it is recommended to prepare aliquots of transformed Agrobacterium directly from the original plasmid stocks. Re-amplification of transformed Agrobacterium cultures is discouraged.

As with the transformation, amplification, and preparation of glycerol stocks in E. coli, it is essential to achieve a high number of colony-forming units (CFUs) when transforming Agrobacterium with *pamiR*. A minimum coverage of 30× (i.e. >70000 CFUs) should be obtained.

The protocol below describes a standard electroporation procedure for electrocompetent Agrobacterium. Depending on the transformation efficiency, multiple parallel transformation reactions may be required. As a starting point, we recommend performing two 100 µl electroporation reactions per amplification.

Supplies needed:

- Agrobacteria with pSOUP helper plasmid. We use the GV3101 strain. Alternatively, the pSOUP helper plasmid can be included into the transformation reaction which might considerably lower the transformation efficiency.
- Electroporation cuvettes
- LB-Medium
- Sterile 2 ml Eppendorf tubes
- Sterile culture container
- Large LB-Agar petri dish supplemented with selection antibiotics (Rifampicin (20 µg/ml), Gentamycin (10 µg/ml) and Kanamycin (50 µg/ml) for GV3103 + pSOUP)→ LB-Agar RBK (from here on)
- 600 ml of liquid LB supplemented with selection antibiotics (Rifampicin (20 µg/ml), Gentamycin (10 µg/ml) and Kanamycin (50 µg/ml) for GV3103 + pSOUP) → LB-RGK in Erlenmeyer flask
- Sterile 60% Glycerol in LB-medium
- Liquid nitrogen

Instructions:

1. All steps should be carried out near a flame or under a sterile hood
2. Thaw electro-competent Agrobacteria on ice
3. Add at least 100 ng but not more than 1.5 µl of plasmids. Make sure the plasmid are stored in very low salt buffer
4. Transfer bacteria and plasmid into electroporation cuvette
5. Apply electric shock (settings for Agrobacteria, one pulse at 2.2 kV)
6. Immediately add 1 ml room temperature LB medium to cuvette, resuspend bacteria and transfer to fresh sterile Eppendorf
7. Recover bacteria by incubating cell suspension at 28°C for 120 min

8. Plate each transformation on two LB-Agar RBK
9. Incubate petri dish at 28°C and check after 1-2 days. You should get very dense but still individual clearly visible colonies. If in doubt count the number of colonies of a fraction of a plate and estimate total CFU
10. Add LB-RGK directly to the plate(s) and re-suspend bacteria using a cell spreader or by pipetting
11. Combine all responded cell suspensions in Erlenmeyer and incubate at 28°C for 180 min
12. Pellet cells from 200 ml by centrifugation (e.g. 2 x 100 ml in 2 x 50 ml falcon tubes) (3500*g* for 15 min). This step can be scaled up if residual culture is not used to directly transform plants.
13. Resuspend pellet(s) in 20 ml cell culture and add 20 ml sterile 60% Glycerol in LB-medium
14. Aliquot into 2 ml Eppendorf tubes and flash freeze in liquid nitrogen
15. Store high density Glycerol stocks at -80°C
16. The residual culture may be used right away to transform the first batch of *Arabidopsis thaliana* plants

#### 1.3 Transformation of the *pamiR* into *Arabidopsis thaliana* plants

Notes:

Use your favorite transformation protocol i.e., whatever gives you the best plant transformation efficiency. However, make sure to always thaw an entire tested glycerol stock to inoculate the culture and select for *pamiR* containing Agrobacteria under antibiotics selection (Rifampicin (20 µg/ml), Gentamycin (10 µg/ml) and Kanamycin (50 µg/ml) for GV3103 + pSOUP). We are using the following protocol:

Supplies needed:

- Sterile culture container (best a 2000 ml baffled Erlenmeyer flask)
- 500 ml of liquid LB supplemented with selection antibiotics (Rifampicin (20 µg/ml), Gentamycin (10 µg/ml) and Kanamycin (50 µg/ml) for GV3103 + pSOUP) → LB-RGK in Erlenmeyer flask
- Floral dip medium: 5% Sucrose and 0.05% Silwet L-77
- *Arabidopsis thaliana* plants

Instructions:

1. Grow 4-5 Arabidopsis plants in one 9 x 9 cm pot until their shoot is approximately 10 cm high
2. Cut this shoot to induce the growth of multiple shoots per plant and wait until every shoot produces flowers (10-15 days)
3. Inoculate 500 ml of LB-RGK in a 2000 ml baffled Erlenmeyer flask with an entire

melted high-density Agrobacteria stock tube

4. Incubate o/N at 28°C with shaking
5. Prepare appropriate volume of floral dip medium
   - We use 500 ml in a 500 ml beaker for dipping
6. Measure $OD_{A600}$
   - Best measure the $OD_{A600}$ in a 1:10 dilution of the culture in LB
7. Calculate the culture volume needed to get a final $OD_{A600} = 0.8$ in anticipated volume used for floral dip
8. Centrifuge cell culture at 3500*g* for 20 min and discard supernatant
9. Resuspend cells in floral dip medium
10. Submerge the flowers of two pots at once in cell suspension for 15 sec. Repeat that step for all plants
11. Subsequently cover plants and store in the dark for 12-16 h
12. Transfer to green house or growth chamber
13. After 10 days, repeat steps 3 to 12 with the same plants

#### 1.4 Selection of transgenic seeds

Notes:

The FastRed seed coat marker is an Olesin-RFP fusion protein (Shimada et al., 2010). This allows for the selection of transgenic seeds by color (see Fig. 1). Transgenic red seeds are visible by eye. It is, however, recommended to select them based on RFP fluorescence. We use a florescence stereo microscope. Spread your seeds on a very smooth surface like a waxed paper or non-static plastic surface and separate red seeds from brown ones. Identification of targeted gene in a selected mutant plant.

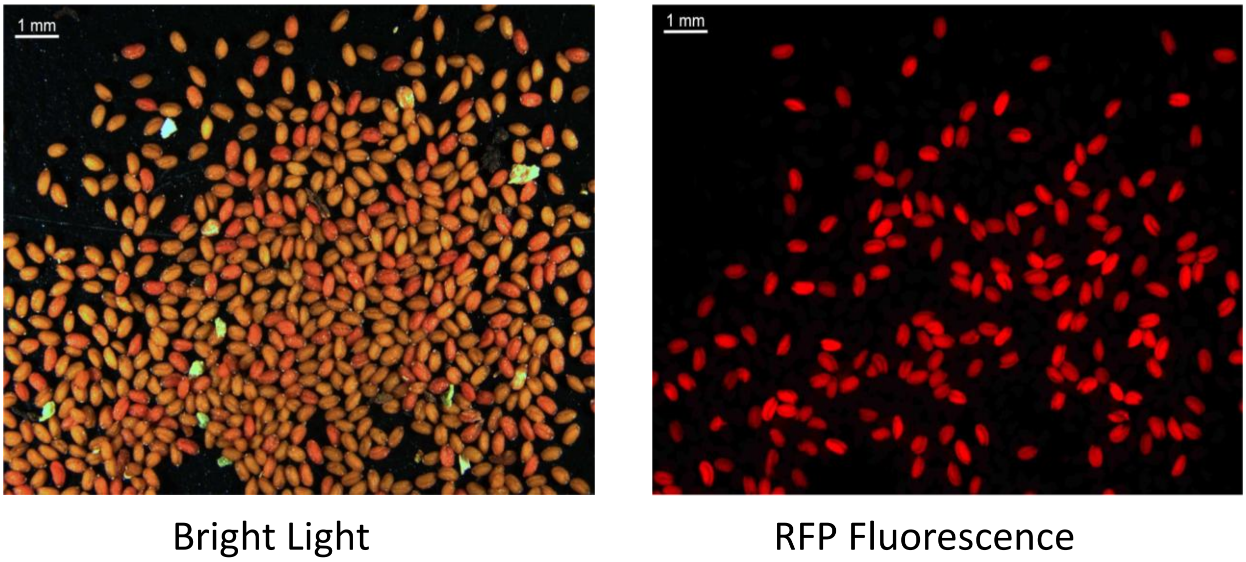


Figure 1: Selection of transgenic seeds using the FastRed seed coat marker. Please note the seed pool in the image is not a $T_1$ pool and parts of these images have been used in Fig.1B of the manuscript. You will see much fewer transformants in $T_1$ (about 1% depending on your transformation protocol).

Notes:

Once you isolated $T_1$ mutant plants from the screen the next step is to identify the causative amiRNA and the targeted gene. This is done by extraction genomic DNA (gDNA), specifically amplifying the amiRNA construct, Sanger sequencing of this PCR product and ultimately using that sequence to identify the amiRNA and the targeted gene.

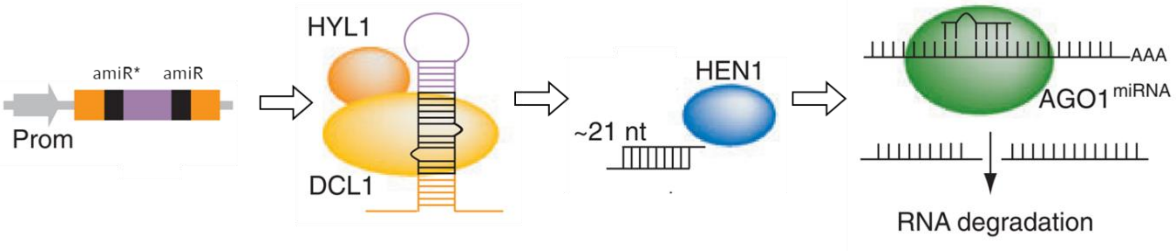


Figure 2: Structure of an amiRNA construct and schematic mechanism of pathway leading to mRNA degaradtion (modified from (Ossowski et al., 2008)).

amiRNA constructs consist of several invariable fragments and two variable sequences. These variable sequences were termed amiR* and amiR sequences. and are responsible for ultimately guiding the AGO1 complex to the targeted mRNA resulting in degradation of the mRNA (see Fig. 2 and (Schwab et al., 2006; Ossowski et al., 2008)). All amiRNAs within the *pamiR* were bioinformatically designed and the targets of all amiR sequences are known. Taken together, this allows for the rapid identification of the targeted mRNA for each amiRNA.

Instructions:

1. Extract gDNA with your favorite method. Low quality gDNA is sufficient. We use a quick and dirty protocol for that (Edwards et al., 1991).
1. PCR-amplify the amiRNA construct using one of the following primer pairs:
   - Primer pair 1: UBQ10_fwd: 5'-GTCGAATAATTACTCTTCGATTTG + mir319_rev: 5'-CCATGGCGATGCCTTA; Product size = 522 bp
   - Primer pair 2: mir319_fwd: 5'- CAAACACACGCTCGGAC + HSP18_rev: 5'-CACAAATTCATAACACAACAAGC; Product size = 517 bp
   - Including a water and a non-transformed gDNA control is recommended.
2. Sanger sequence the PCR product using any of the primer used for the amplification
   - We are directly sequencing 1 µl of the PCR product without prior purification
3. Align sequencing result with the *pamiR* example vector sequence section.
   - A .gb file of the annotated example vector in which the amiR* und amiR sequences are marked is provided. This will look like this (Fig.3):

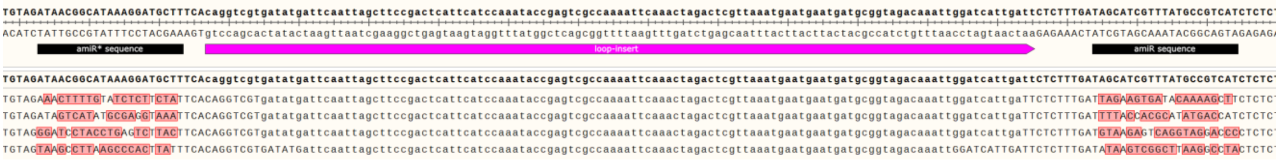

Figure 3: Example for an alignment of the Sanger sequencing result of an amiRNA candidate with the example vector. Image taken from the Snapgene software

4. Copy the amiR or amiR* sequence of the Sanger sequencing and go to the provided

excel sheet “Supplementary Table S1-*pamiR* ami sequences and targeted genes.xlsx”. Search the document using the copied sequence.

5. The AGI(s) of the targeted gene(s) will be in the column D.

Final remarks:

The amiRNAs were designed using the Phantom database (Hauser et al., 2013). The web page has discontinued. Using the amiR sequence you can also do a reverse search to check for targeted genes on the homepage of the Weigel lab (http://wmd3.weigelworld.org/cgi-bin/webapp.cgi?page=TargetSearch;project=stdwmd).

#### 1.5 How to handle candidates

Notes:

As with all large-scale screening methods, candidates have to be confirmed. There is a multitude of possibilities which could lead to phenotypes independent of the amiRNA, such as insertion of the cassettes into existing genes or regulatory elements, genome alterations due to t-DNA insertions, activation of genes by the UBQ10 promoter of the amiRNA, etc. However, based on our experience, these effects happen much less often than phenotypes caused by amiRNA-mediated knock down of gene transcripts.

We recommend doing some of following steps to confirm candidates identified with above mentioned methods:

- Back-cross your candidate with the original plant line (e.g., Col-0 wild type). Use the same conditions than used in the screen and see whether the phenotype penetrates in the $F_1$. Keep in mind that a proportion of your transgenic seeds might not be as efficient in silencing the target genes than the $T_1$ plant.
- Check $T_2$ seed segregation by simply checking the RFP fluorescence of your seeds.
- Check the phenotype of the transgenic $T_2$ seeds.
- Clone the same amiRNA which is causing the initial phenotype, transform your original plant line and check transgenic $T_1$ seeds for the phenotype.
- Order t-DNA insertion lines or use CRISPR-CAS9 to create mutants in the target gene.
